# Robust representation and non-linear spectral integration of simple and complex harmonic sounds in layers 4 and 2/3 of primary auditory cortex

**DOI:** 10.1101/2025.08.26.672221

**Authors:** Yunru Chen, Chih-Ting Chen, Yuhan Gui, Patrick O. Kanold

## Abstract

Sound harmonicity is foundational in complex auditory stimuli like music and vocalizations but it remains unclear how such spectrally complex stimuli are processed in the auditory cortex (ACtx). Subregions of the auditory cortex process are thought to process harmonic stimuli differently, and secondary ACtx (A2) layer (L) 2/3 is believed to be the most selective. Selective responses to sound features in ACtx are thought to emerge hierarchically starting from A1 L4. Since the spectral complexity of harmonic stacks can range from two to more than ten components, harmonic selectivity and sensitivity might also arise hierarchically across layers and areas. We studied responses to simple and complex harmonic stacks across A1 L4, A1 L2/3, and A2 L2/3 in adult mice using in vivo two-photon microscopy. We found harmonic-sensitive neurons (HN) responding only to harmonic stacks but not to their pure-tone components in all areas at similar proportions. HNs showed non-linear processing of component tones with onset-responsive HNs showing greater nonlinearity, which decreased with harmonic complexity. Co-tuned HNs in A1 L4 exhibited the highest signal correlation, regardless of harmonic complexity. A1 L4 HNs also showed the lowest noise correlation with other neurons. Moreover, A1 L4 HNs achieve robust spectral integration and harmonic sensitivity by receiving diverse inputs and maintaining high signal correlation, ensuring independent, strong responses to harmonic stimuli. Therefore, harmonic sensitivity is present in A1 L4 and is not a unique feature of A2. Thus, tuning to complex spectral sounds is a fundamental property of ACtx and is already established in A1 L4.

**Significance statement:** Harmonics are essential in auditory perception, influencing how we process complex sounds like music and speech. This study reveals that neurons in the primary auditory cortex (A1) and secondary auditory cortex (A2) integrate simple and complex harmonic structures with distinct mechanisms of neuronal recruitment. A1 L4 harmonic-sensitive neurons (HNs) demonstrated strong, independent responses through high signal correlation and minimal noise correlation, suggesting a robust mechanism for spectral integration. Our results show that harmonic relationships are already extracted at the input layers of A1, and that HNs show non-linear facilitative integration. Thus, tuning to sounds of complex spectral contents might be a fundamental processing function of the auditory cortex and is already established in A1 L4, which receives major thalamic inputs.

## Introduction

Harmonicity is a fundamental feature of human speech, music and animal vocalization (Ehret and Riecke, 2002). Harmonic stacks are compound sounds made up of one or more frequencies that have a positive integer multiple of the fundamental frequency (F_0_) (Terhardt, 1974; Plack, 2010; McLachlan et al., 2013). Both simple sounds such as pure tones and sweeps and complex sounds like animal vocalizations are represented in the cochlea and, more sparsely, in the central nervous system (Nelken, 2004; Hromadka et al., 2008; Chechik and Nelken, 2012). In mice, ultrasonic and middle frequency vocalization usually consist of a few frequency components, whereas vocalization in low frequency harmonics is composed of multiple frequencies. Behavioral studies reveal that a high proportion of low frequency harmonic vocalizations are associated with stress in mice (Grimsley et al., 2013; Grimsley et al., 2016) and serve as a distress call to listeners (Chen et al., 2009). As a central hub for sound processing, the auditory cortex (ACtx) encodes the spectral and temporal information in harmonic stacks to form a neural representation of presented auditory stimuli (Zatorre and Belin, 2001; Kadia and Wang, 2003; Linden et al., 2003; Bendor et al., 2012; Herdener et al., 2013). Studies in animal models have shown that the ACtx represents various sound types-such as sweeps, white noise, and pure tones-through sparse coding (Hromadka et al., 2008; Liang et al., 2019; Kang and Kanold, 2024). Specifically, neurons that are selective to harmonic stacks have been identified in both the core and belt areas in ACtx of ferrets (Bizley et al., 2010), the primary auditory cortex (A1) of cats (Abeles and Goldstein, 1970; Phillips and Irvine, 1981; Sutter and Schreiner, 1991), in A1 of marmosets (Kadia and Wang, 2003; Bendor and Wang, 2005; Song et al., 2016). In mouse A1, L2/3 and L4 display distinct responses of pure tones and unique integration patterns for the spectral content of harmonic stacks (Sołyga and Barkat, 2022). Others suggested that A2 L2/3 shows a preferential response to harmonics than individual pure tone components compared to A1 and anterior auditory field (AAF) (Kline et al., 2021). Thus, harmonic processing might vary between ACtx subareas and layers. The mammalian brain contains two hemispheres and in mice, imaging has shown that A2 in the left hemisphere of mice exhibits a stronger response to high-frequency tones and vocalizations than the right hemisphere, analogous to human left-hemisphere specialization for speech (Calhoun et al., 2023), suggesting that harmonic processing might also vary between hemispheres.

Furthermore, studies suggest that neurons may integrate simpler and more complex harmonic stacks differently. In marmosets, neurons in A1 have been shown to respond to more complex harmonic structures selectively, exhibiting non-linear facilitation compared to responses to pure tones or two-tone harmonic stacks (Song et al., 2016; Feng and Wang, 2017). This sensitivity is further underscored by the disruption of neuronal activation when the spectral content of perfect-octave harmonic stacks is shifted, indicating high sensitivity for precise spectral composition. While sparse coding in the cortex is well-documented, it remains unclear how this coding differs for simple versus complex harmonic stacks. Specifically, it is unknown whether a fundamental neural network recruits additional neurons or strengthens neuronal connections to process more complex harmonics that are composed of simpler harmonics.

The mammalian ACtx is organized into layers. In rodents, layer (L) 4 (L4) neurons receive primary thalamic inputs (Winer et al., 2005; Sherman et al., 2013; Ji et al., 2016), while L2/3 neurons receive inputs from both L4 and other L2/3 neurons within auditory cortical subregions through horizontal connections (Oviedo et al., 2010; Covic and Sherman, 2011; Watkins et al., 2014; Meng et al., 2017; Chang and Kanold, 2021). Since inputs and outputs of each ACtx subarea are unique (Hackett, 2011), harmonic stimuli might be processed differentially by specialized neuronal populations distributed across layers in both primary and higher-order auditory fields.

While the spectral integration of complex sounds in A1 L2/3 and L4 has been investigated using electrophysiology (Sołyga and Barkat, 2022), there is a lack of a direct comparison of how selectively tuned neuron populations across layers and subregions are recruited and interconnected to process the same harmonic stacks. Moreover, it remains an open question whether the integration of complex harmonic stacks builds upon the integration of simpler harmonic stacks, and whether this integration occurs hierarchically or in parallel across A1 L4, A1 L2/3, and A2 L2/3.

In this study, we aimed to determine (i) whether harmonics of varying spectral complexity are represented in similarly sparse manner across A1 L4, A1 L2/3, and A2 L2/3, (ii) whether harmonic-sensitive neurons encode harmonic stacks with comparable consistency across these subregions, and (iii) whether functional connectivity between harmonic-sensitive neurons and other sound-responsive neurons differs among three populations. We analyzed sound-evoked activity in mouse left ACtx subregions using in vivo two-photon Ca^2+^-imaging to capture the simultaneous activity of large neuron populations. We used awake adult CBAxThy1GCaMP6s F1 mice that have good high-frequency hearing (Frisina et al., 2011; Bowen et al., 2020) and express the calcium indicator GCaMP6s in excitatory neurons. We find that harmonic-sensitive neurons (HNs) are present in similar proportions in A1 L4, A1 L2/3, and A2 L2/3, regardless of harmonic complexity. However, co-tuned HNs in A1 L4 displayed the highest functional connectivity, the highest signal correlation, and the lowest noise correlation compared to A1 L2/3 and A2 L2/3. We also observed nonlinear integration of the harmonic component frequencies. The integration of component frequencies became more linear with increasing spectral complexity. These results suggest that while HNs maintain similar sparseness across cortical subregions, A1 L4 uniquely supports efficient, robust integration of spectral contents through strong co-tuning and minimal noise, enabling consistent encoding of both simple and complex harmonic structures. Thus, harmonic stimuli are processed in all auditory subareas, starting with the input layers of ACtx, suggesting that harmonic processing might be a core function of ACtx.

## Methods

### Animal Preparation

For in vivo two photon imaging, mice were 12-24 weeks old at the time of the experiments. Mice (n=28) of both sexes were first generation (F1) of CBA/CaJ (Jax # 000654) and C57BL/6J-Tg(Thy1-GCaMP6s)GP4.3Dkim/J (Jax # 024275) crosses. For electrophysiology recording, mice were Tlx3-Cre;Rosa26-LSL-GCaMP6s;cdh23^Ahl/ahl^ mice on a C57BL/6J background with fixed-hearing (Babola et al., 2025) All mice were housed with a reverse light cycle (12h light, 12h darkness). All experiments were conducted during the dark cycle of the mice. All animal procedures were approved by the Johns Hopkins University Animal Care and Use Committee.

### Surgery

In brief, mice were prepared for surgery by inducing anesthesia with 4% isoflurane in 100% oxygen and later reduced to 1-2% isoflurane for maintenance until the surgery ended. Dexamethasone (2mg/mL) was injected 1 hour before the surgery started to prevent inflammation. During the surgery, body temperature was maintained with a feedback-controlled heating pad maintained at 35-36°C. Hair on the top of the head was shaved and removed using a hair remover (Nair) followed by disinfections with 70% ethanol and iodine. The skin and soft tissues were then exposed by detaching and pushing away muscle on the surface of the skull. Craniotomy of a circular area with 4mm diameter was performed above the left ACtx, covering both A1 and A2, by using a dental drill. A stack of round glass coverslips (one 4mm glass, catalog #640724-CS4R, Warner Instruments, on two stacked 3mm glasses, catalog #64-0720-CS-3R, Warner Instruments, and fixed with optic glue, catalog #NOA71, Norland Products) was secured onto the exposed brain with SuperGlue around the edge of the window. The exposed skull was covered with dental cement (C&B Metabond). To prepare mice for imaging, a customized metal headplate was fixed onto the cement along the midline of the skull. After the surgery, 5 mg/kg carprofen and 500 mg/kg cefazolin were injected subcutaneously after the surgery and in the following 1-2 days of surgery. Mice rested in the home cage and recovered for at least 14 days before the first imaging session. The surgical procedure for acute electrophysiological experiments was similar to the imaging experiments described above, except that the circular craniotomy was done with dura removed. The exposed brain was covered by a 4mm glass with saline filling the gap. We used silicone adhesive (KWIK-SIL, World Precision Instruments) to seal the cranial window for the ease of removal before the experiment. Cefazolin, dexamethasone, and meloxicam (5mg/mL) were injected after surgery, and the animal was put in a recovery cage for at least one hour before the experiment started.

### Sound Stimuli

All sound stimuli were pre-generated using MATLAB (Mathworks, version R2023a). Stimuli were loaded into a Tucker-Davis Technologies (TDT) RX6 processor and presented by a ES1 speaker, via a PA5 attenuator, 10 cm away from the right ear of the mouse. The speaker was first calibrated using customized MATLAB (Mathworks, version R2020b) scripts by recording 4– 64 kHz pure tone frequencies at 70 decibels (dB) sound pressure level (SPL), with a calibrated microphone to find the natural transfer function of the speaker. We then calculated the inverse of the function, which when added to the natural transfer function of the speaker, will equalize the speaker’s output, giving a flat frequency/dB curve. We then used the calibration curve to generate each pure tone frequency at 70 dB SPL and attenuated each frequency to 60 dB SPL. We ensured that the recorded sound level of each frequency component was <1 dB from the target for all sounds played using MATLAB (Mathworks, version R2023a). To study the effect of the complexity of the spectral contents in a harmonic complex, we generated harmonic complexes that are composed of two to ten frequencies with base frequency 4kHz as well as complexes that are composed of two to five frequencies with base frequency 8kHz. Similar to multiple published studies that focus on the encoding of complex sounds in mice (Wang et al., 2020; Kline et al., 2021; Sołyga and Barkat, 2022; Kang and Kanold, 2024; Efron et al., 2025), we chose this frequency range because sounds composed of low-to-mid frequencies have been shown in mouse vocalizations, pup calls, and alarm calls in mice, which makes the encoding of these sounds relevant and potentially critical to social behaviors (Ehret and Bernecker, 1986; Ehret and Riecke, 2002; Grimsley et al., 2013; Grimsley et al., 2016). To generate harmonic stacks with varied spectral complexities, each frequency component was generated at 60dB SPL and stacked to generate the harmonic stacks without further attenuation. By doing this, each frequency component during presentations of harmonic stacks or pure tones has the same sound intensity. The sound intensity of resulting harmonic stacks can vary from 63 dB (two-tones harmonic) to 70dB (ten-tones harmonic). For spectrally shifted two-tone harmonic stacks (SH), we generated each frequency component similarly and stacked two 60dB SPL frequency components, so the sound intensity of the resulting two-tone JH was 63dB SPL, comparable to two-tone harmonic stacks. During two-photon imaging, each session started with a 10-second of silence period to record the spontaneous activity of neurons. Each trial comprises of 0.5 s of pre-stimulus silence, 1 s of sound presentation, followed by 3.5 s post-stimulus silence (Fig. 1A). The order of sound presentation, including harmonic stacks and pure tones, is pseudo-randomized to ensure all stimuli are played once before beginning the next randomized sequence. Each sound was repeated for 10 times to increase statistical confidence.

**Figure 1:**
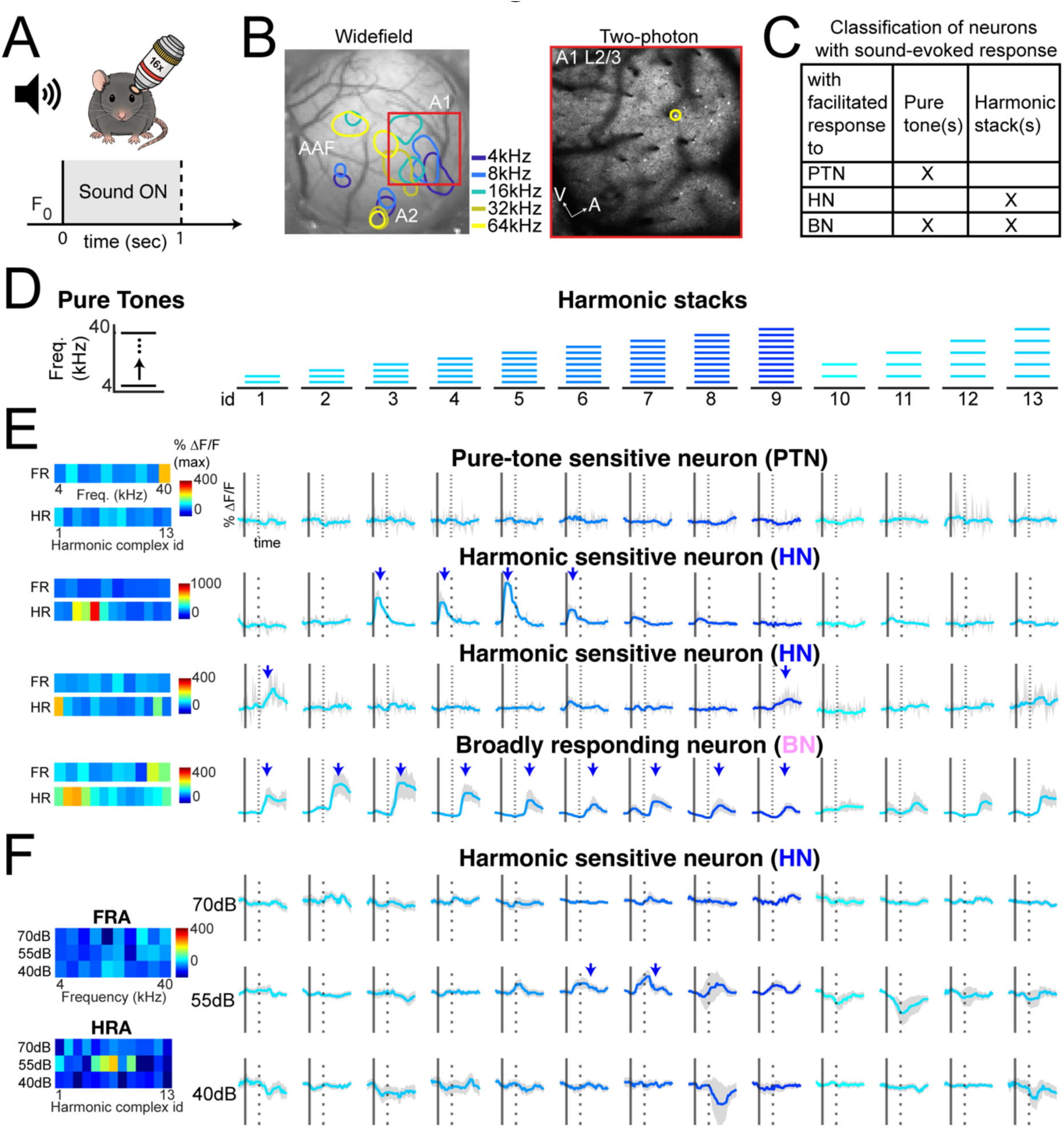
Excitatory neurons in A1 L4, A1 L2/3, and A2 L2/3 are found sensitive to harmonic stacks but not to individual frequencies. **A:** In vivo two-photon imaging schematic (top) and sound stimulus trial (bottom). F_0_: baseline of fluorescence change in the window of neuron response to the sound stimulus. Solid line indicates sound onset and dashed line indicates sound offset. The gray shaded area indicates when sound is present. **B:** Left: Tonotopy map (A1, A2, AAF) identified by widefield imaging on sound-evoked areas. Right: Example field of view of in vivo two photon calcium imaging on A1 L2/3 excitatory neurons. **C**: A decision chart of classifying neuron into three categories based on their facilitative response to types of sounds. **D**: Schematic of stimuli design. Pure tone frequency, for example the base frequencies 4kHz and 8kHz, is shown in black. Harmonic stacks that consist of more than one frequencies are shown in blue. **E:** Examples of fluorescence changes compared to baseline (ΔF/F) of neurons responding to only pure tones (PTN), only to the harmonic stacks (HN) with onset or offset response, or both harmonic(s) and pure tone(s) (BN) (from top to bottom). For HN and BN, their ΔF/F responses to the pure tone frequency components are stacked and shown as black traces as the leftmost plot. **F**: Example of one HN responding to pure tones and harmonic stacks played at three sound levels, 70dB (top row), 55dB (middle row), and 40dB (bottom row). Arrows indicate detected significant facilitated response to the corresponding stimulus.

### In vivo two-photon calcium imaging

The location of the ACtx and subareas, including A1, A2, and AAF, was acquired by the characteristic tonotopic axes using widefield imaging (Liu et al., 2019). During wide field imaging, mice were head-fixed on a customized imaging station under the 2x objective. Mice were presented with 100ms 4kHz to 64kHz pure tone frequencies in three sound levels (60-, 75-, and 90-dB SPL). For two-photon imaging, mice were head-fixed on a customized imaging station under the 16x objective attached to the two-photon microscope within a sound-attenuating chamber. In one imaging session, harmonic stacks and the stack components were randomized and played as described above. GCaMP6s was excited at 940 nm, and the field of view contained 1024 × 1024 pixels covering 1120 x 1120 μm^2^. Images were acquired from either L2/3 (150-230 um below the surface) or L4 (370-430 μm below the surface) using the Prairie View software at 15 frames per second. Data from each subarea was acquired in separate sessions. For recording in A2 L2/3, 1.1x-1.5x zoom is used to restrict the field of view to contain only the tonotopy-mapped A2. Sound stimulation was synchronized with the image acquisition using a hardware trigger signal.

### Two-photon data analysis

Motion correction and cell extraction were performed using the suite2p software with denoising (Pachitariu et al., 2017). Neuron fluorescence traces and neuropil fluorescence traces were extracted by processing recorded signals with Suite2P software. Pixels that overlapping more than one cell were excluded from processing. Denoising was enabled for data processing in Suite2P. Neuron fluorescence were further corrected as: F_cell, corrected_ = F_cell_ – 0.9 * F_neuropil_. We calculated the change of fluorescence (**Δ**F/F) during response period by dividing fluorescence 3 seconds after the sound onset from each trial by the average fluorescence of silent frames in 0.5 seconds preceding the sound onset (F0). To determine that a neuron has facilitated response to a sound, we calculated the confidence for F_0_ and **Δ**F/F in all trials of one sound, respectively, and set our criterion to be that the lower bound of **Δ**F/F must be larger than the upper bound of F0 at 5% confidence level. The suppressed response is not considered in this study.

### Classification of neurons by sound-evoked response

To categorize neurons into harmonic-, pure tone-, or both-sensitive neurons, we compared their sound-evoked response calculated as described above. Neurons with no significant facilitative response to any pure tone but with significant response to any harmonic stacks were categorized as harmonic-sensitive neurons (HNs). Similarly, those with no significant facilitative response to any harmonic stacks but with significant facilitative response to any pure tone sounds were categorized as pure tone-sensitive neurons (PTNs). Those with significant facilitative response to both harmonic stacks and pure tone sounds were considered as both-sensitive neurons (BNs).

### Linearity analysis

To evaluate the linearity and nonlinearity of harmonic neurons responding to harmonic stacks, we used customized MATLAB scripts to perform univariate and multivariate linear regressions as well as Support Vector Regression for nonlinear regression on the mean response to pure tone frequencies and harmonic stacks. For univariate linear regression, we used the MATLAB function “polyfit” with degree of 1 to linearly reconstruct harmonic response, y_n_, by adjusting the coefficient assigned to each pure tone response X separately. n represents reconstructed harmonic response by response to pure tone n. We reported the highest R2 values of all univariate regressions for one harmonic response.

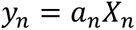

For multivariate linear regression, we used MATLAB function “regress” to perform linear reconstruction by response to all corresponding pure tone components.

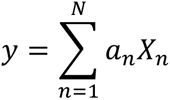

To evaluate the effects of varied number of predictors on the multivariate regression, we permuted predictor data, or response to pure tone, then performed multivariate regression again and compared the R^2^ between original data and permuted data. To perform Support Vector Regression analysis, we used the MATLAB function “fitrsvm” with the radial basis function (“rbs”) kernel and data standardization.

### Granger causality analysis

We investigated functional connectivity by performing Granger causality analysis on the ΔF/F traces of HNs and BNs. We used the multivariate Granger causality framework implemented in the MVGC toolbox (Barnett and Seth, 2014). For neuron in this analysis, we extracted all trials for all sounds and concatenate all traces into one time-series data, which were then detrended and z-scored. To reduce the effect of slow fluctuations, we applied a first-order difference filter to each time-series data. GC was then estimated using vector autoregressive (VAR) models fit to each pair of neuron traces between every HN and every BN within each FOV. The optimal model order was selected to be 1 due to the slow calcium dynamics.

Statistical significance of GC links between one HN and any BN within the same FOV was determined by comparison to a nonparametric null distribution generated through bootstrapped surrogate data. For each BN-HN pair, we generated 5 surrogate traces by permutating frames of the BN’s time series data, totally disrupting the temporal dependency between the pair. GC values from real data were considered significant if they exceeded the 95th percentile of the surrogate distribution (p < 0.05, one-tailed). To obtain ΔGC values, we subtracted the real GC value of each BN-HN pair by the mean of GC values of surrogate data.

All GC values were calculated across the entire stimulus period (or spontaneous window, where relevant), and results were analyzed at the individual neuron-pair level and aggregated to quantify the distribution of ΔGC strength across sound conditions and subareas. Only neuron pairs with sufficient activity (above threshold signal variance and number of time points) were included in the final analysis. The distance between BN-HN is normalized by dividing the absolute distance by the maximum value of distances of all BN-HN pairs within the same FOV. To perform Spearman correlation analysis, we used the built-in MATLAB function “corr” with type of correlation being “Spearman”.

### Onset-offset bias index (OBI)

We investigated the onset-offset bias index using the same methods described in our previous study (Liu et al., 2019). We defined the onset response window to be the 0.5 seconds immediately following sound onset, and the pre-onset baseline window to be 0.5 seconds before the sound onset. Similarly, we defined the offset response window to be 0.5 seconds immediately following the sound offset, and the baseline for the offset response to be 0.5 seconds before the sound offset. For each neuron, we first averaged the ΔF/F of ten trials for each window, then calculated the onset response as the mean activity in the onset window minus the mean of the pre-onset window, and the offset response as the mean activity in the offset window minus the mean of the pre-offset window. These values were then used to compute the OBI as:

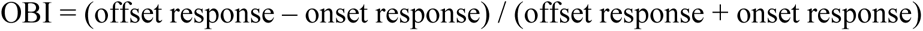

### Electrophysiology experiment and data preprocessing

Extracellular recordings were performed with a 4-shank high-density probe (Neuropixels 2.0) (Steinmetz et al., 2021)in an anechoic chamber, where the animal was head-fixed and awake during the recording. Audio stimuli were presented with a free-field speaker (ES1, TDT) positioned 10cm from the animal’s right ear, contralateral to the recording hemisphere. The speaker was driven by a pre-amplifier (ED1, TDT) receiving input from a data acquisition device (NI USB-6343). Neuropixels recordings were acquired using the SpikeGLX software system, and the data was digitized at 30kHz. The neural recording system and the sound stimulation were coordinated by custom scripts in MATLAB. For spike analysis, a global demuxed CAR (common average referencing) filter and a bandpass filter (300 Hz-9000 Hz) were applied to correct for temporal misalignment across channels due to multiplexing during analog-to-digital conversion at the electrode sites and remove irrelevant signals using post-processing tool CatGT. Spike sorting was done by Kilosort4 (Pachitariu, 2023) , and we only included those clusters that were classified as single units in the analysis. Current source density (CSD) was derived from the second derivative of local field potential (LFP) (Mitzdorf and Singer, 1978)with a spacing of 200*μ*m. The LFPs were obtained from applying a bandpass filter at lower frequencies (0.1Hz-500Hz) to the recordings of all channels. We used CSDs to localize the relative laminar position of each recording site. L4 was characterized by a short latency sink after the stimulus onset, whereas the peaks of current sources marked L2/3 and L5.

### Analysis of electrophysiology data

To determine if the single unit was responsive to the stimuli, we counted the spike number before and after the stimulus onset for each trial and used paired t-test to determine whether the spike count within 0.2s after the sound onset is significantly different from the spike count within 0.2s before the sound onset. Similarly, to account for the offset response, we used a paired t-test to determine whether the spike count within 0.2 before the sound offset is significantly different (p < 0.001) from the spike count within 0.2 after the sound offset. Clusters with significant onset or offset response with increased spike counts were then further classified into three categories: PTN, HN, and BN based on whether the clusters had evoked facilitated response to pure tones and/or harmonic stacks.

### Correlation analysis

To compute signal correlations, the pair-wise cross-correlation of sound-evoked activity was calculated between co-tuned neurons using 2 seconds after the sound onset, covering the offset period, similar to our previously published studies (Winkowski and Kanold, 2013). To compute the pair-wise noise correlations between each neuron pair, ΔF/F of each trial of a sound is subtracted by the mean ΔF/F of all trials of the sound, then the cross correlation between the time sequences was calculated for each trial between paired neurons with significant sound-evoked facilitated response by using the MATLAB function “xcorr” at zero lag and normalized by using “coeff” as the normalization parameter.

### Statistical analysis

All statistical tests were performed in MATLAB R2024b. The Anderson-Darling test was used on each data group to test for normal distribution. For data that does not form normal distribution, a two-sided Wilcoxon rank sum test or a Kruskal-Wallis test was performed and corrected by Bonferroni correction for statistics obtained by multiple comparisons Dunn’s test followed by Dunn-Sidak multiple comparison correction. For data that follows normal distribution, an N-way Analysis of Variance (ANOVA) test is performed followed by post-hoc t-test and corrected by Tukey-Kramer.

## Results

To explore the neural representation of harmonic stacks across three auditory subregions (A1 L4, A1 L2/3, and A2 L2/3), we first used widefield imaging to identify tonotopic maps and localize A1 and A2 in each mouse (Fig. 1B) as described previously (Liu et al., 2019). We then performed in vivo two-photon imaging in A1 L4, A1 L2/3, and A2 L2/3 in separate sessions (3 sessions per animal) on awake young adult mice (A1 L4: n=11; A1 L2/3: n=10; A2 L2/3: n=10 animals). We used F1 mice with a CBA background, which retain high-frequency hearing into adulthood (Frisina et al., 2011; Bowen et al., 2020), to ensure responses to the high sound frequencies in complex harmonic stacks. We presented either pure tones (PT) or harmonic stimuli (H) with varied numbers of components (2–10 components), covering frequencies from 4–40 kHz at 60–70 dB SPL. Harmonic complexes were centered on fundamental frequencies (F0) of 4 kHz and 8 kHz (Fig. 1C).

### HNs exhibit high sensitivity for harmonic tuning at one sound level

The ACtx is thought to have a sparse representation of sound features, so we first investigated whether a subset of excitatory neurons responded exclusively to harmonic stimuli without responding to any pure-tone components of the harmonic stacks at one sound level and whether different subfields displayed a varying fraction of harmonically responsive neurons. We recorded sound-evoked responses to harmonic stimuli with varied component numbers, as well as to individual pure tones (A1 L4: 10 animals, 4974 neurons; A1 L2/3: 12 animals, 10,913 neurons; A2 L2/3: 7 animals, 1792 neurons). Imaging sessions in A1 L4, A1 L2/3, and A2 L2/3 were conducted when the targeted subarea was clearly identified in widefield imaging and suitable for two-photon imaging. Each frequency was presented at the same sound intensity to ensure the same intensity of each component of one harmonic sound compared to when each frequency is played at pure tone stimulus.

We classified neurons into three types based on their facilitated (positive ΔF/F) responses to sounds: pure-tone neurons (PTNs; responsive only to pure tones), harmonic neurons (HNs; responsive only to harmonic stimuli), and broadly-tuned neurons (BNs; responsive to both pure tones and harmonic stacks) (Fig. 1D). All three subregions exhibited similar proportions of HNs, PTNs, and BNs (Fig. 4A-C), suggesting comparable sparseness in harmonic encoding across these areas. We further quantified the proportion of neurons activated by different levels of harmonic complexity in each individual recording session, and found that the number of neurons with facilitated responses remained largely consistent between areas as the harmonic stacks became more complex (Fig. 4). Male and female mice had the same proportions of sound-evoked neurons (Fig. 4A-C); therefore, data from both sexes were combined in further analyses. Even though we did not observe an effect of increasing the spectral complexity of harmonic stacks, we found that the proportion of neurons activated by harmonic stacks sound in A2 L2/3 was higher than those in A1 (Fig. 4D, left). When we considered the tuning properties, A2 L2/3 also contained a larger fraction of BNs than A1 L4 and L2/3 across a range of complexities (Fig. 4D, middle), while A1 L4 contained a smaller fraction of HNs than A1 L2/3 and A2 L2/3 (Fig. 4D, right). These results indicated that all subareas contain a similar proportion of neurons responsive to only the harmonic stacks, but that A2 contains more neurons responsive to pure tones and harmonic stacks (BNs).

The proportion of activated neurons reflects the network size representing harmonic stacks, while response amplitudes of individual neurons may reflect the robustness of the sound representation by the neuronal network, which is crucial for stable perception. We thus calculated the response amplitude of each HN to its “best harmonic“—the harmonic stimulus eliciting the strongest average response. Our results revealed that HNs in A1 L2/3 showed significantly higher response amplitudes to their best harmonic stacks compared to HNs in A1 L4 (Fig. 4E). Together, these findings suggest that neurons in all three subregions exhibit sensitive tuning to harmonic stacks, with HNs in A1 L2/3 showing the highest activation by harmonic stacks. Additionally, our results also show that the sensitivity is observed at the thalamorecipient layer (A1 L4), suggesting that the source of selectivity could reflect both cortical processing and thalamic inputs. However, the higher response amplitudes in A1 L2/3 suggest a more robust representation of the harmonic stacks in this layer compared to A1 L4. We observed no differences in amplitude between A2 L2/3 and the other two subareas, which suggests that A2 L2/3 HNs are more heterogeneous in terms of response amplitude to the same sounds that are represented significantly differently in A1 L2/3 and A1 L4.

To identify the sensitivity of HN neurons to harmonic stacks as opposed to sounds having multiple frequency components that are not harmonically related, we performed separate imaging sessions presenting two-tone harmonic stacks and non-harmonic two-tone sounds with the upper frequency component spectrally shifted downward (SHs) (Fig. 5A). We found that in all three subareas fewer HNs responded to both the harmonic and its spectrally shifted counterpart (Fig. 5B). We observed no differences in the proportion of neurons responsive to 25%, 50%, and 75% spectrally shifted harmonic stacks across the three subareas (Fig. 5C). These results suggest that a subset of excitatory neurons in three imaged subregions have sound-evoked facilitative response only to the specific frequency combination forming a harmonic stack and are sensitive to the downward spectral shifts of the higher frequency in a two-tone harmonic stack.

Our findings show that HNs display high selectivity to harmonic stacks, and the representation of more complex harmonic stacks does not rely on recruiting additional HNs that respond to simpler harmonic stacks, as shown by similar proportions of neurons evoked by harmonic stacks of varied complexity. This further supports the hypothesis that ACtx neurons are finely tuned to spectral information within harmonic stacks and represent these features in a sparse and selective manner.

### Electrophysiology recording of HN response to harmonic stacks in both L4 and L2/3

Two-photon imaging in deeper layers has the potential of contamination by neuropil signal from more superficial layers. Thus, to confirm our observation about harmonic-sensitive neurons in L4 by two-photon imaging, we performed electrophysiology recording in awake hearing-fixed mice mice on a C57BL/6J background (Babola et al., 2025). To record across the depth of A1 we used Neuropixel arrays (REF) and identified cortical layers using CSD analysis (Fig. 2B). We identified recording locations in L4 as sites showing the first current sink shortly after sound onset as L4 and sites 200 µm above the early current sink as L2/3. We then identified clusters with significant sound response within 0.2 after onset or offset and found clusters that show similar sensitivity to harmonic sounds (HNs) but not to pure tones (Fig. 2C). We found HNs in both A1 L2/3 and L4 (Fig. 2D). This result is consistent with our in vivo two photon imaging results (Fig. 1) and suggest the existence of neurons that are responsive selectively to harmonic stacks but not to any individual pure tones in both L4 and L2/3 of A1.

**Fig. 2:**
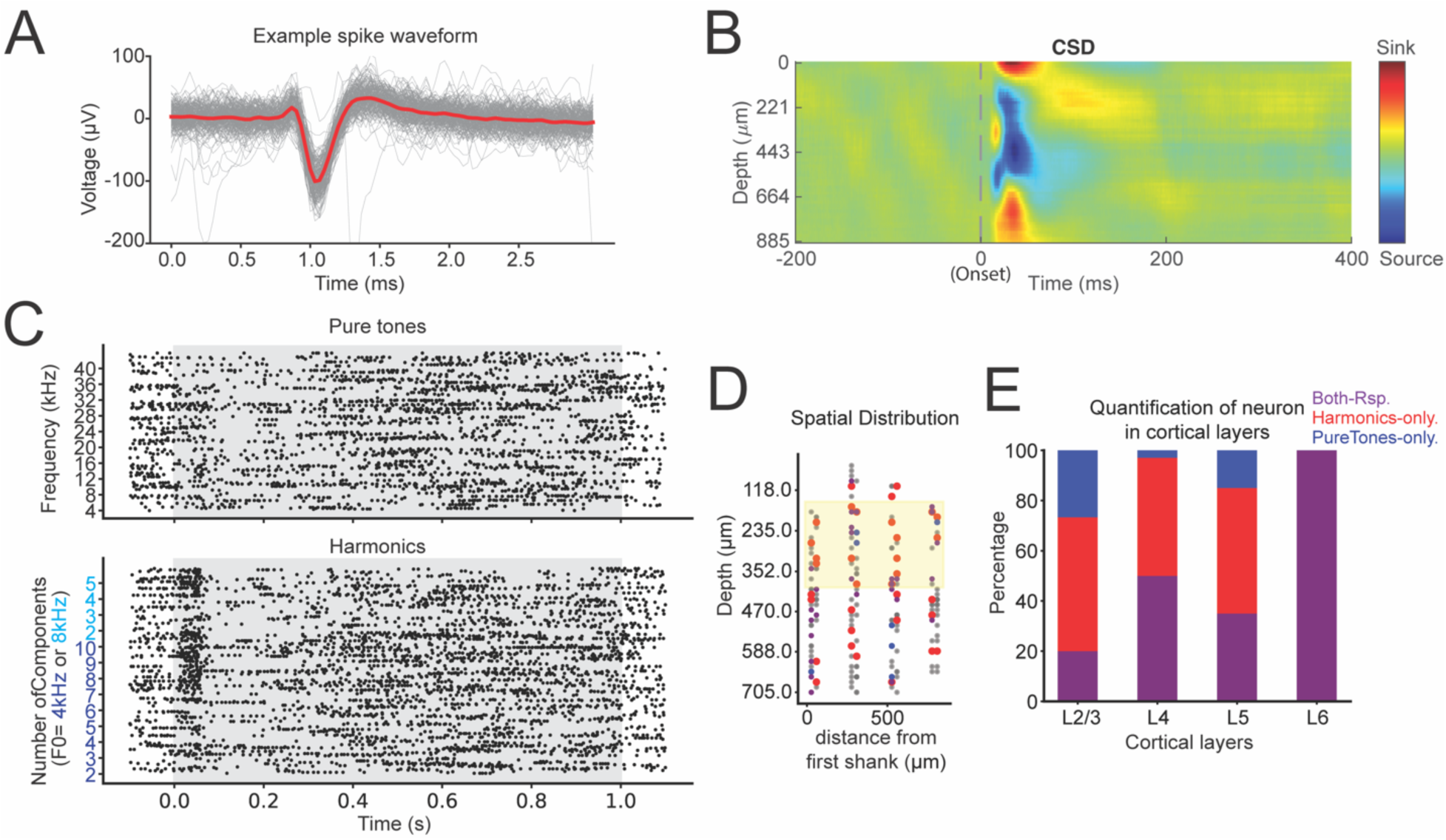
Electrophysiology recording confirms existence of harmonic-sensitive neurons in ACtx. **A: Example waveform of spikes from the electrophysiology recording. Grey lines show the 200 example traces aligned based on spikes. Red line show the average of the 200 aligned traces.** **B**: Example plot of current source density (CSD) analysis for identifying L4 which shows current sink after sound onset, and L2/3 which shows current source and is located above L4. **C**: Example raster plots of a recorded cluster that show significantly increased counts of spikes within 0.2s window after the sound onsets of harmonic stacks (bottom) and non-significantly changed spike counts after sound onset. Gray area indicates the presence of sound (1s). **D**: Distribution of clusters in a representative mouse that show significantly increased spike counts to pure tones and/or harmonic stacks. Same criteria for classification as in vivo two-photon imaging is applied to classify clusters as HN (red), PTN(blue), or BN(purple). Gray are clusters that have no significant sound-evoked response to any sound. Yellow shaded area indicates L4. **E**: Quantification of proportions of classified clusters in different cortical layers of the same mouse in (**D)**.

### BN shows broader frequency response profiles compared to PTN

In our previous work (Liu and Kanold, 2021; Maximov and Kanold, 2025), we identified six distinct shapes of frequency response area (FRA) of neurons that respond to pure tones of different frequencies at different sound levels. We here again studied the FRAs of the neurons responding to pure tones and/or harmonic stacks as a separate experiment. We imaged A1 L2/3 and A1 L4 in hearing-fixed mice and played pure tones and harmonic stacks at three sound levels, 40dB, 55dB and 70dB. For harmonic stacks, each frequency is calibrated to 40dB, 55dB or 70dB then combined to generate the harmonic stacks. We found the best frequency for each BN and PTN and used the best frequency as the center of the FRA. We then clustered the centered FRA of each BN and PTN similarly, as in our previous work (Liu and Kanold, 2021), into one of the clusters V, H, I, S1, S2, S3 (Fig. 3D). We found that tSNE plot exhibited distinct clusters (Fig. 3A). BN and PTN occupied different regions of the tSNE plots (Fig. 3B) which is further analyzed by quantifying the proportions of BN and PTN in each cluster (Fig. 3C). We found that proportions of PTNs in S1 and S3 clusters are significantly higher than the proportions of BNs (Fig. 3C). For cluster S2, proportion of PTNs showed a trend to be higher but not significant compared to the proportion of BN. This result indicates that PTNs usually respond to their best frequency at one level and have much narrower tuning regarding pure tones compared to BNs. Additionally, for cluster H, proportion of PTNs showed a trend to be lower but not significant compared to the proportion of BNs. This result further suggests that, compared to PTNs, BNs tend to have broader tuning reflected by FRA shape and they respond to more neighboring frequencies around the best frequencies when the sound becomes louder.

**Fig. 3:**
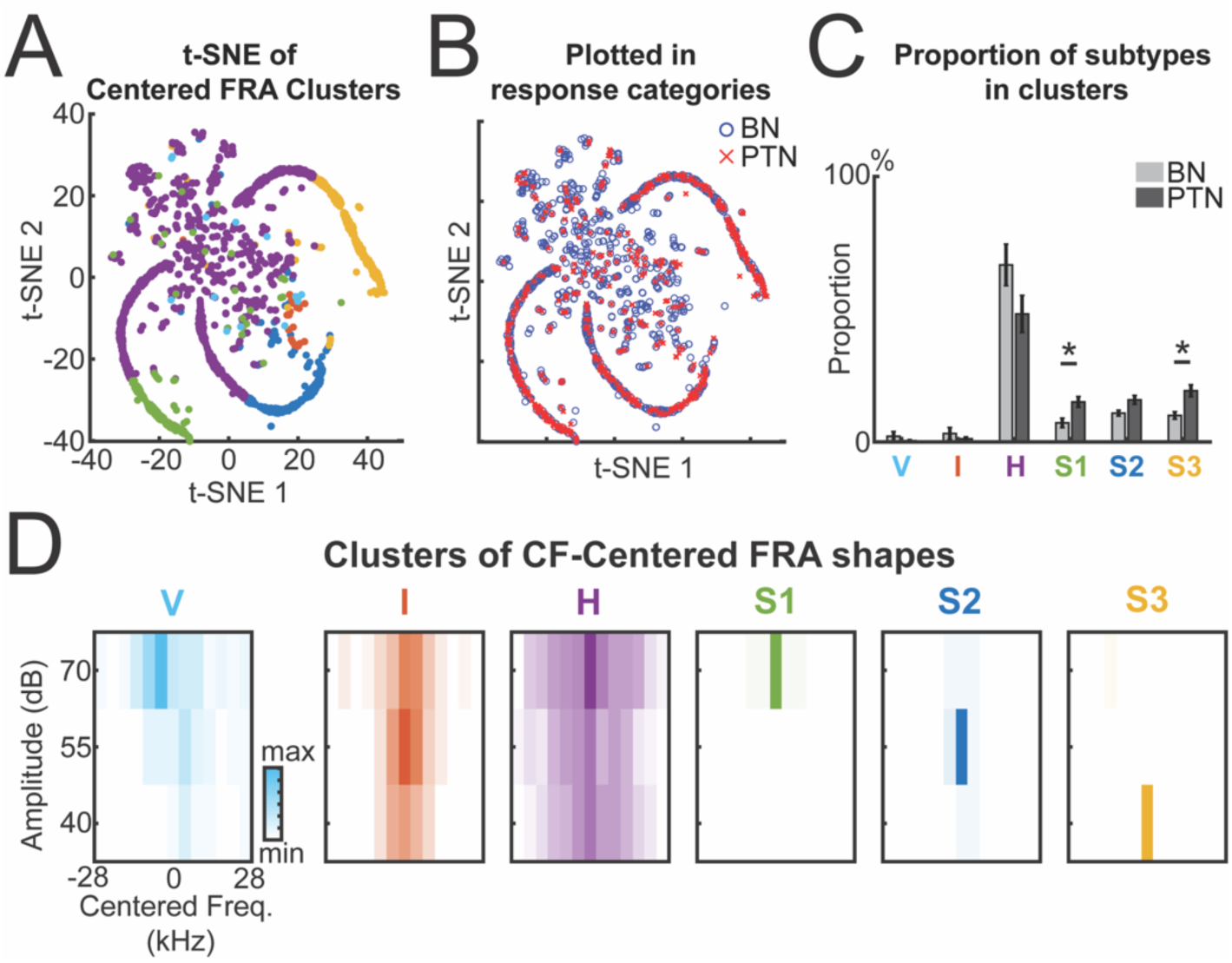
Distinct frequency response area (FRA) profiles of broadly tuned neurons (BNs) and pure-tone neurons (PTNs) in A1 L2/3. **A**: t-SNE visualization of k-means clustering applied to centered FRA profiles from sound-responsive neurons (n = 3 animals, 1418 neurons). **B**: Same t-SNE plot as in (A), now colored by BN and PTN identity, illustrating the distribution of neuron subtypes across FRA-based clusters. **C**: Proportion of BNs and PTNs within each cluster, showing differential cluster occupancy by neuron subtype. Error bars indicates standard errors. *: p < 0.05. Two-sample t-test on proportions of BN versus PTN for each cluster: V, p = 0.4125; I, p = 0.4543; H, p = 0.1531; S1, p = 0.0445; S2, p = 0.0680; S3, p = 0.0249. **D**: Average centered FRA maps for each of the six clusters, revealing distinct spectral and intensity tuning profiles. Vertical bars indicate peak response frequency.

**Figure 4:**
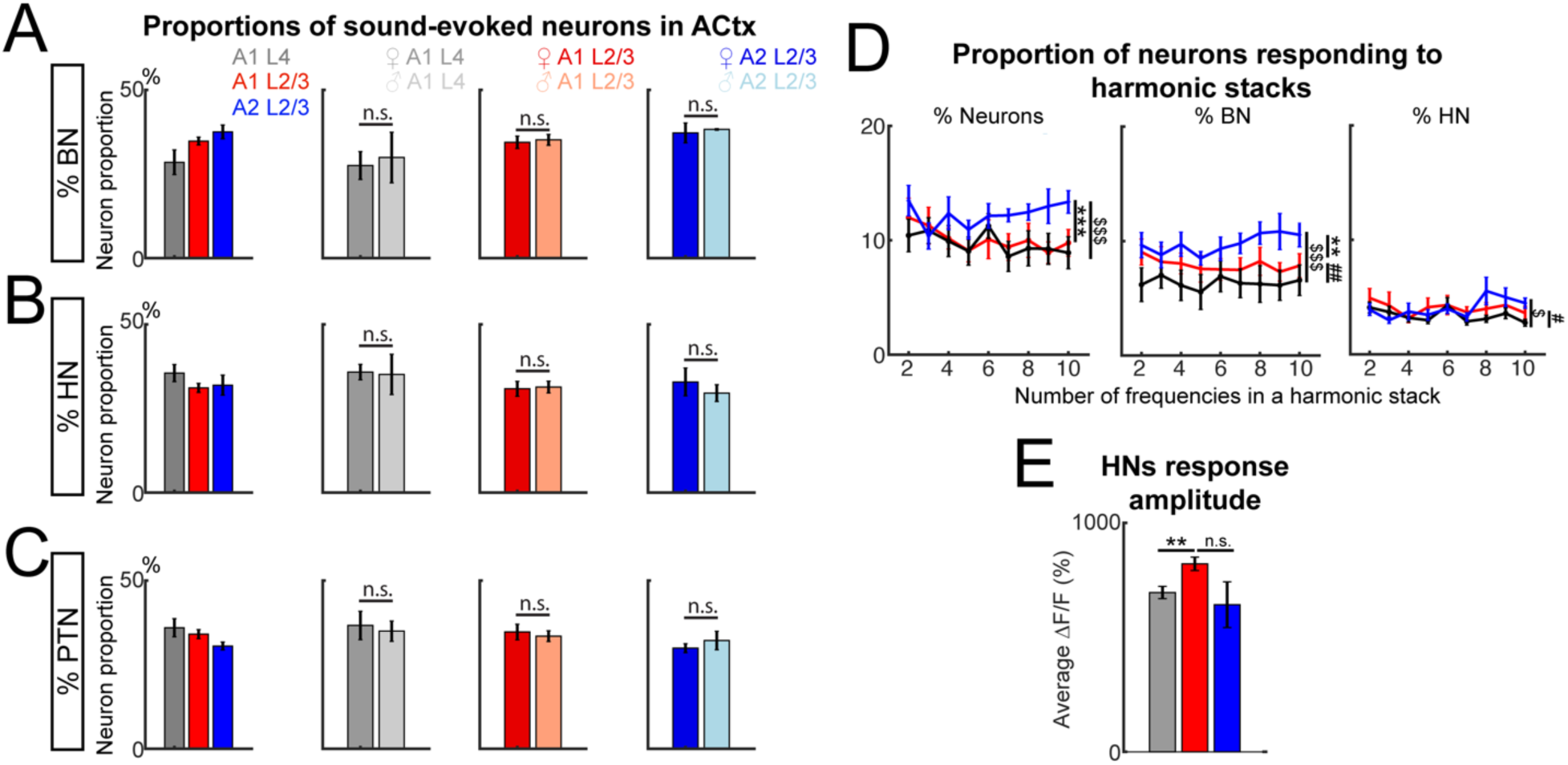
Harmonic neurons are similarly sparse in three imaged auditory subareas. **A:** Left: Proportions of BNs in A1 L4, A1 L2/3 and A2 L2/3. **%BN**: A1 L4, 28.50%±11.60%; A1 L2/3, 34.80% ± 4.09%; A2 L2/3, 37.52% ± 5.42%. One-way ANOVA on the effect of subregions: F(2,26) = 3.17, p = 0.059. Middle and right: comparisons of BN proportions in females and males in subregions. A1 L4: Female, 27.54%±10.17%; Male, 29.92%±15.05%. One-way ANOVA on the effect of sex: F(1,8) = 0.0907, p = 0.77. A1 L2/3: Female, 34.44%±4.56%; Male, 35.17%±3.96%. One-way ANOVA on the effect of sex: F(1,10) = 0.0877, p = 0.7732. A2 L2/3: Female, 37.22%±6.6%; Male, 38.26%±0.34%. One-way ANOVA on the effect of sex: F(1,5) = 0.0443, p = 0.8416. Error bars show mean ± SEM. **B**: similar to (**A)**, but proportions of HNs. **%HN**: A1 L4, 35.54%±8.09%; A1 L2/3, 31.12% ± 4.68%; A2 L2/3, 31.94% ± 7.83%. One-way ANOVA on the effect of subregions: F(2,26) = 1.24, p = 0.307. A1 L4: Female, 35.81%±5.62%; Male, 35.12%±11.97%. One-way ANOVA on the effect of sex: F(1,8) = 0.0157, p = 0.9035. A1 L2/3: Female, 30.86%±5.44%; Male, 31.37%±4.3%. One-way ANOVA on the effect of sex: F(1,10) = 0.0326, p = 0.8604. A2 L2/3: Female, 32.89%±9.22%; Male, 29.58%±3.54%. One-way ANOVA on the effect of sex: F(1,5) = 0.2211, p = 6580. **C**: similar to (**A)**, but proportions of PTNs. **%PTN**: A1 L4, 35.97%±8.49%; A1 L2/3, 34.08% ± 4.70%; A2 L2/3, 30.54% ± 3.12%. One-way ANOVA on the effect of subregions: F(2,26) = 1.67, p = 0.21. A1 L4: Female, 36.64%±10.31%; Male, 34.96%±6.06%. One-way ANOVA on the effect of sex: F(1,8) = 0.0850, p = 0.7780. A1 L2/3: Female, 34.70%±5.68%; Male, 33.46%±3.93%. One-way ANOVA on the effect of sex: F(1,10) = 0.1907, p = 0.6692. A2 L2/3: Female, 29.89%±3.01%; Male, 32.15%±3.88%. One-way ANOVA on the effect of sex: F(1,5) = 0.7131, p = 0.4369. **D:** Proportions of neurons responding to harmonic stacks with varied spectral complexity in A1 L4 (left), A1 L2/3 (middle), and A2 L2/3 (right). Two-way ANOVA: %neurons: Subareas, F(2,234) = 11.5, p = 1.72E-5; component numbers, F(8,234) = 1, p = 0.43; interactions between subareas and component numbers, F(16,234) = 0.61, p = 0.87. %BN: Subareas, F(2,234) = 19.52, p = 1.44E-8; component numbers, F(8,234) = 0.31, p = 0.96; interactions, F(16, 234) = 0.31, p = 0.99. %HN: Subareas, F(2,234) = 4.99, p = 0.0076; component numbers, F(8,234) = 1.97, p = 0.051; interactions, F(16,234) = 1.21, p = 0.26. Post-hoc t-test with corrected p-values on subareas: %neurons: A1 L4 v.s. A1 L2/3, p = 0.75; A1 L2/3 v.s. A2 L2/3, p = 1.67E-4; A1 L4 v.s. A2 L2/3, p = 1.84E-5. %BN: A1 L4 v.s. A1 L2/3, p = 0.0028; A1 L2/3 v.s. A2 L2/3, p = 0.0014; A1 L4 v.s. A2 L2/3, p = 1.33E-9. %HN: A1 L4 v.s. A1 L2/3, p = 0.013; A1 L2/3 v.s. A2 L2/3, p = 0.99; A1 L4 v.s. A2 L2/3, p = 0.029. All data are represented as mean ± SEM. Symbols indicate significance with post-hoc t-test with Bonferroni correction. *: p<0.05, **: p<0.01, ***: p<0.001. * indicates comparison between A1 L2/3 and A2 L2/3. # indicates comparison between A1 L2/3 and A1 L4. $ indicates comparison between A1 L4 and A2 L2/3. **E:** Response amplitude of HNs responding to the characteristic harmonic sound. Error bars show mean ± SEM. A1 L4: 6.95 ± 0.27; A1 L2/3: 8.21 ± 0.29; A2 L2/3: 6.42 ± 1.00. Wilcoxon rank-sum test with Bonferroni-corrected p-values: A1 L4 v.s. A1 L2/3: p=0.012; A1 L2/3 v.s. A2 L2/3: p = 0.36; A1 L4 v.s. A2 L2/3: p = 0.056.

**Figure. 5:**
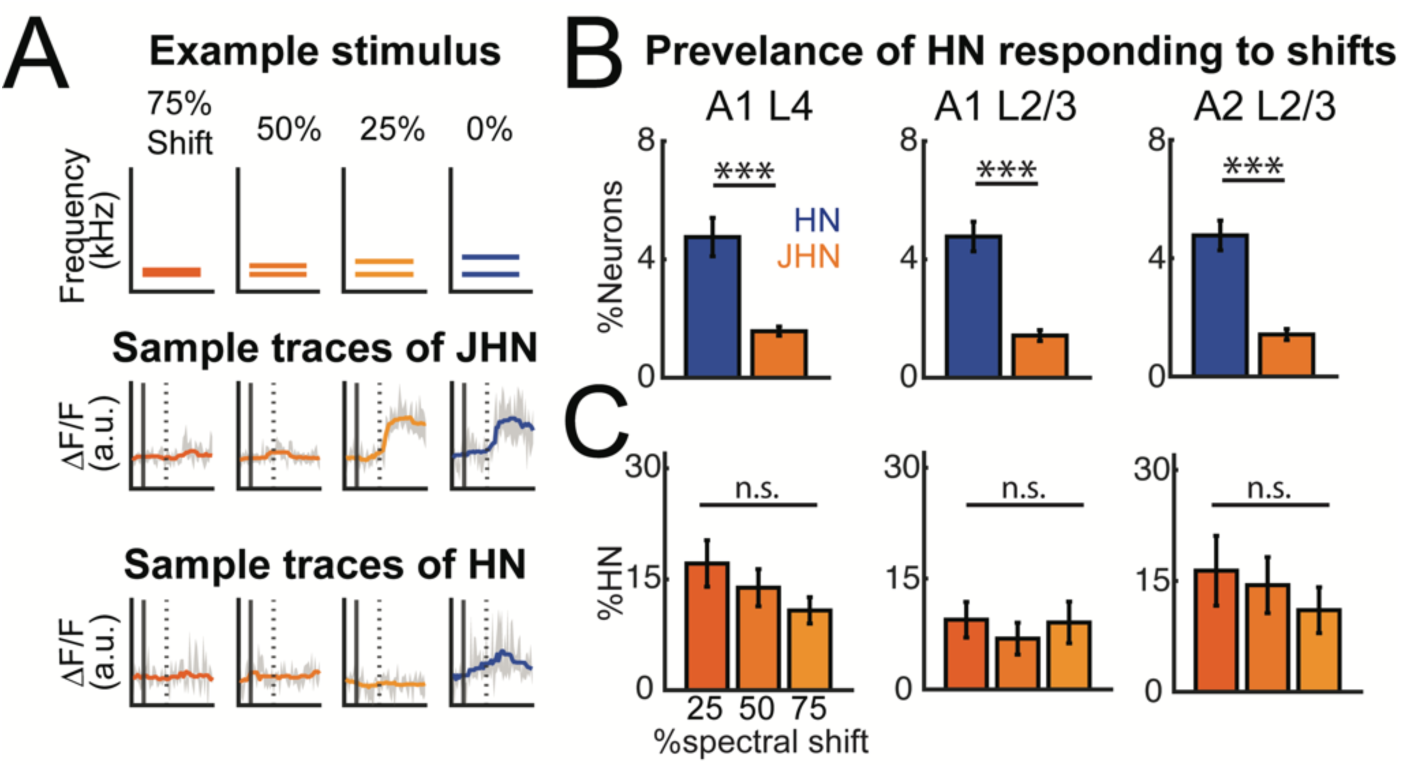
Most of HNs are sensitive to frequency shifts in harmonic stacks and faithfully respond to the harmonic stacks. **A**: Example of one two-tone harmonic and its spectrally shifted stacks (top) and example of ΔF/F of neurons only responding to the harmonic (blue) but not spectral shifts (orange), as well as shift-tolerant harmonic neurons (SHN) with facilitated response to both the harmonic and its spectral shifted versions but not to any pure tone frequencies that makes up the composite sounds. **B**: Prevalence of HNs and SHNs in three subareas. One-way ANOVA: A1 L4 (7 mice), F(1,16) = 22.74, p = 4.58E-4; A1 L2/3 (6 mice), F(1,16) = 111.56, p = 9.61E-7; A2 L2/3 (9 mice), F(1,16) = 39.01, p = 1.17E-5. **C**: Distribution of SHN across spectral shifts within the HN population in A1 L4, A1 L2/3 and A2 L2/3. One-way ANOVA: A1 L4 (7 mice), F(2,16) = 1.56, p = 0.24; A1 L2/3 (6 mice), F(2,16) = 0.31, p = 0.74; A2 L2/3 (9 mice), F(2,16) = 0.47, p = 0.63.

Together with our criteria to classify neurons as BNs in Figure 1, these results further suggest that BNs may contribute more to the encoding of spectral combinations and may serve as an intermediate population bridging basic and complex spectral features, whereas PTNs serve in precise frequency resolution at one sound level.

### Nonlinear and linear integration of pure-tone components in harmonic sensitive neurons

How does the sensitivity to harmonic stacks emerge? Harmonic responses can occur via linear or non-linear processing of the individual sound components. Given that we identified HN neurons in all areas, these processes might vary between areas. We examined if the responses of HN neurons to harmonic stacks could be due to processes beyond the simple summation of responses to each pure-tone component, which suggests a potential nonlinearity that might vary with increasing spectral complexity of harmonic stimuli. We thus investigated whether HN responses to harmonic stacks were constructed linearly or non-linearly from the individual pure-tone components. We classified harmonic responses as linear if they could be accurately approximated by a weighted sum of these components as a first-order polynomial function (Fig. 6A). Conversely, if the sum could not fully account for the harmonic response, we inferred the presence of nonlinear processing.

**Figure 6:**
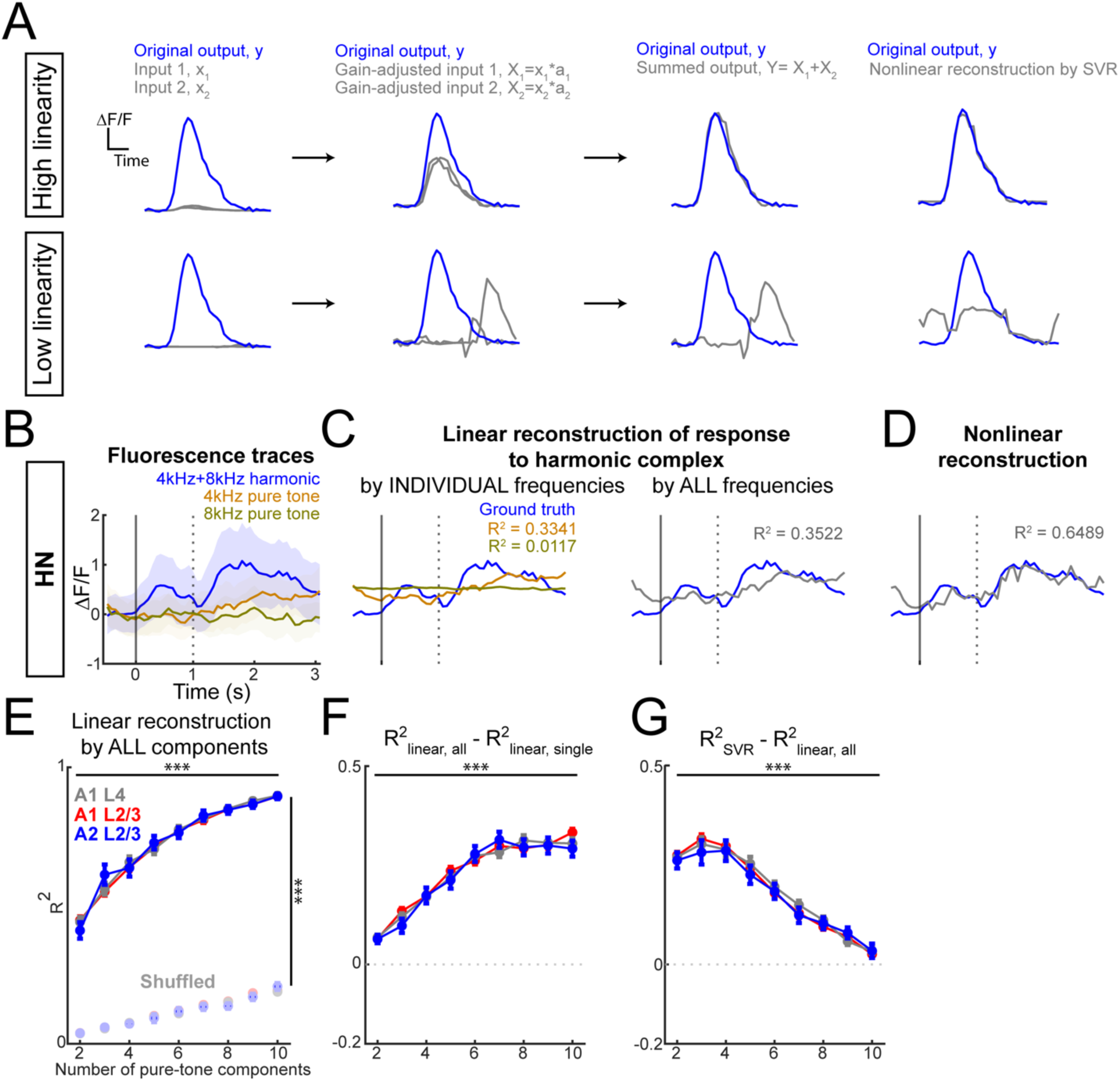
Non-linearity of response of HNs diminishes when harmonic becomes more spectrally complex. **A**: Schematic showing the linear reconstruction of the simulated output signals by taking the sum of gain-adjusted simulated input signals. Cases of high linearity (top) and low linearity (bottom) are shown. **B**: Example response to harmonic stacks and pure-tone components of a HN with best harmonic being 4kHz + 8kHz stack. **C**: Linear reconstruction of the example HN response to harmonic by using both the response to the pure tones 4kHz and 8kHz as variables (left), as well as by the response to each pure tone separately (right). Higher R^2^ tells higher similarity between reconstructed harmonic response by component response and the original response to harmonic. **D**: Nonlinear reconstruction with Support Vector Regression (SVR) that accounts for nonlinearity in the relationship between three responses, which shows higher R^2^ score. **E**: Quantification on the similarity between reconstructed response to harmonic and original response in A1 L4, A1 L2/3, and A2 L2/3. Left: Similarity by using linear reconstruction improves as the harmonic becomes more spectrally complex, which is not shown in permuted response data while the same number of independent variables are used in the reconstruction. Three-way ANOVA (factors: subareas, harmonic stacks, permutation): Subareas, F(2,6406) = 0.0058, p = 0.9942; harmonic stacks, F(8,6406) = 372.44, p = 0; permutation, F(1, 6406) = 24375.9, p = 0; Subareas x Harmonics, F(16,6406) = 0.52, p = 0.9404; Subareas x Permutation, F(2,6406) = 0.71, p = 0.49; Harmonics x Permutation, F(8,6406) = 147.0647, p = 5.8806E-228. **F**: The similarity between reconstructed response and original response when the reconstruction uses response to all components as variables versus response to single component. Two-way ANOVA (factors: subareas, harmonic stacks): Subareas, F(2, 3195) = 0.7822, p = 0.4575; harmonic stacks, F(8, 3195) =167.9672, p = 0; interaction, F(16,3195) = 0.9670, p = 0.4907. **G**: Right: The similarity between reconstructed response and original response when SVR that accounts for more non-linearity in the data was used compared to linear reconstruction. Two-way ANOVA (factors: subareas, harmonic stacks): Subareas, F(2, 3195) = 0.54, p = 0.5822; harmonic stacks, F(8, 3195) =111.86, p = 0; interaction, F(16,3195) = 0.43, p = 0.9763.

To quantify linearity, we first isolated HN responses to their optimal harmonic stimuli and their constituent pure-tone components (Fig. 6B). Using these responses, we reconstructed the HN response to harmonic stacks by summing the pure-tone responses through first-order polynomial fitting (Fig. 6C). Additionally, we employed support vector regression (SVR) to capture further nonlinear relationships between harmonic and pure-tone responses (Fig. 6D). To assess the linearity, we calculated the R² as a measure of the goodness of fit for each polynomial function. To control for the number of predictors in the multivariate function’s effect on R², we permuted values across the averaged neuronal responses and conducted the same linear reconstruction, allowing us to perform a three-way ANOVA to compare R² across spectral complexity, cortical subareas (A1 L4, A1 L2/3, and A2 L2/3), and data type (original versus permuted) (Fig. 6E). Our ANOVA revealed significant effects for spectral complexity, data type, and their interaction. However, no significant differences were found between subareas, suggesting that HNs in these regions integrate individual frequencies similarly across simple and complex harmonic stacks.

The analysis further revealed that R² values increased with harmonic complexity, indicating that more responses to spectrally complex harmonic stacks are due to a more linear computation to integrate individual frequency components. We then examined whether this linear reconstruction was driven by responses to one or multiple frequency components by comparing R² values from single-component and multi-component reconstructions (Fig. 6F). For more complex harmonic stacks, all-components reconstructions yielded significantly higher R² values, implying that the response integration for complex harmonic stacks leverages the contributions of multiple frequency components more linearly.

To probe further into the nonlinear aspects of harmonic processing, we applied SVR to account for nonlinearities in HN responses. We observed improved R² with SVR relative to linear reconstruction. However, the degree of improvement diminished as harmonic stacks became more complex (Fig. 6G), supporting our finding that HNs exhibit greater linearity with increasing spectral complexity. This heightened linearity in complex harmonic stacks may be due to reduced inhibitory modulation in HNs, while nonlinearity might stem from two primary sources: (i) threshold effects due to intrinsic properties (e.g., ion channel density, leaky conductance, and inhibitory inputs) (Gerstner et al., 2014), and (ii) amplification effects from positive feedback mechanisms that exponentially boost the pure-tone response into a harmonic response (Wu and Zenke, 2021). In conclusion, our data suggest that HNs processing complex harmonic stacks might experience reduced inhibition and positive feedback compared to those processing spectrally simpler harmonic stacks (Fig. 6E-G). To summarize these effects, we propose a simple model microcircuit (Fig. 6H) in which an excitatory HN tuned to two-tone harmonic stacks receives both inhibitory and excitatory inputs from PTNs. When only one pure tone is present, the HN is non-responsive due to balanced excitation from a single PTN and inhibition. However, when both tones are presented as a two-tone harmonic, the HN’s response is facilitated by simultaneous excitation from both PTNs and the disinhibition effect, leading to a nonlinear, harmonic-sensitive response. Given that all subareas show similarities in the HNs we propose that such a circuit is present across ACtx.

### A1 L4 shows the highest BN-HN directed functional connectivity

As we revealed that the response of HN to harmonic stacks cannot be reconstructed by a linear weighted sum of the neuron’s response to pure tone components, we performed the same analysis for BNs. This analysis showed that the reconstructed response of BN to harmonic stacks is more linear (Fig. 7A-C). Combined with its broad tuning, BN might play a critical role in enabling selective response to harmonic stacks in HNs. Here, we proposed a micronetwork to explain the source of harmonic-sensitive tuning of HNs as excitatory neurons (Fig. 7D). We focused on investigating the potential differences of functional connectivity (FC) from BN to HN by applying Granger Causality (GC) analysis on the sound-evoked response of BNs and HNs. We aimed to answer two questions: first, does BN-HN have significant FC compared to the shuffled control? Second, does the FC differ between subareas and across distances?

**Fig. 7:**
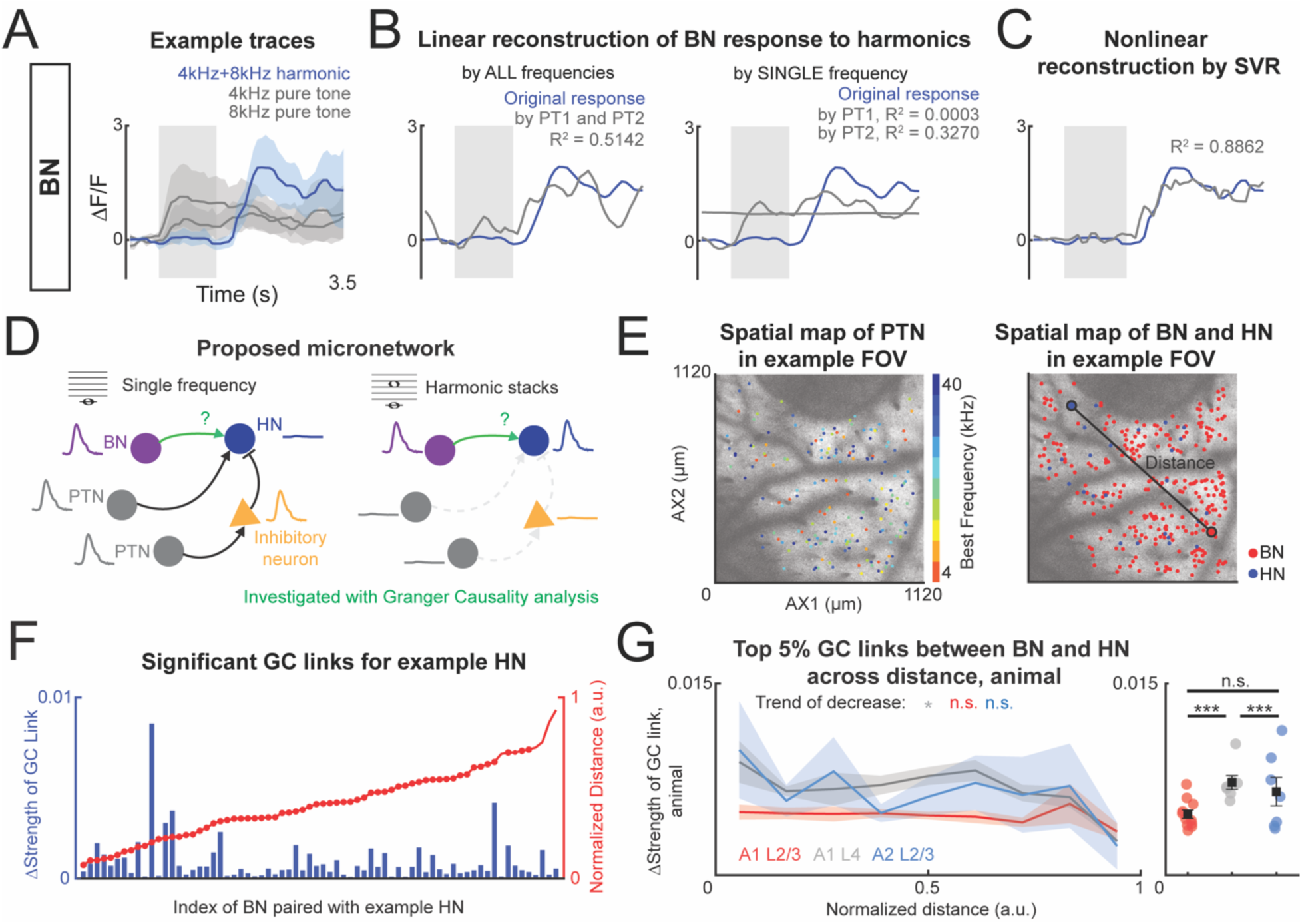
BN-HN in A1 L4 has the highest functional connectivity. **A:** Example realistic neural responses of BNs at different pure-tone sounds and corresponding harmonic sounds. Case of harmonic response (4 kHz plus 8 kHz) and pure-tone responses (4 kHz and 8 kHz) are shown. **B:** Linear reconstruction of the example BN response to harmonic sound by using both the response to the pure tones as variables (left), as well as by the response to each pure tone separately (right). Higher R^2^ indicates higher similarity between reconstructed harmonic response by component response and the original response to harmonic. **C:** Nonlinear reconstruction of BN responses to HN response by Support Vector Regression (SVR). **D:** Schematic showing the proposed micronetwork that can encode how HN responses to sounds with different components. Along the auditory pathway, PTN activates inhibitory neurons when pure-tone sound is played, further inhibits the HN (left). BN directly activates BN without any inhibition from PTN, which triggers the high responses of HN when harmonic is played. We next investigated the functional connectivity between BN and HN by applying Granger Causality (GC) analysis. **E:** Example field-of-view images showing the spatial location of PTNs, HNs and BNs in A1 L2/3. Left: PTNs are located heterogeneously across imaged A1 L2/3. Color indicates best frequency of the plotted neuron. Right: BN and HN are plotted on the same field-of-view. Larger dots indicate an example coactivated pair of HN and BN to one stimulus, in which the BN achieves the highest R^2^ reconstructing the response of HN to the stimulus. The black line indicates the distance between the HN and BN. **F:** ΔStrengths of significant GC links between all BNs within the same FOV (left Y-axis, blue) as an example HN plotted with the distances between each BN and the example HN (right Y-axis, red). The distances are normalized by the maximum distance among all BN-HN pairs for all HNs for each animal. **G**: Quantification of top 5% GC links between any BN-HN pair plotted against binned normalized distances in three subareas. A1 L2/3, n = 11 mice. A1 L4, n = 7 mice. A2 L2/3, n = 7 mice. Two separate Kruskal-Wallis tests revealed the significant main effect of subareas (χ²(2) = 35.3033, p = 2.16E-8) and non-significant effect of distances (χ²(8) = 13.2486, p = 0.1036) on the Δstrength of top 5% GC links. Post-hoc Dunn’s test with Dunn-Sidak correction showed that A1 L4 has the highest functionally connected BN-HN pairs compared to A1 L2/3 (p = 1.03E-8) and A2 L2/3 (p = 5.71E-4). Spearman correlation performed on the GC links as the distances increase reveals significant trend of decrease of GC links in A1 L2/3 (ρ = -0.2170, p = 0.0388) and nonsignificance in A1 L4 (ρ = -0.1202, p = 0.3731) and A2 L2/3 (ρ = -0.2132, p = 0.1254).

To answer these questions, we performed the Granger causality analysis on every pair of BN-HN to explore their functional connectivity. We used the metric, ΔGC link strength, to quantify the relative strength of GC links by subtracting absolute GC link values of the real data by the mean of absolute GC link values of the surrogate data. By doing this, we extracted the strength of GC links in real data compared to chance, which would still generate values of GC links. To answer the first question of significant GC link between BN and HN, we found that more than one BN-HN pair for each mouse showed significant GC links (Fig. 7F), despite the absolute values of GC links being small (<0.05), which can be due to the slow calcium dynamics or sparse functional coupling. We then answered the second question by performing statistic tests on two main factors: subareas and distances. We found no significant effect of distances between BN and HN on the strength of GC links (Fig. 7G). However, the test revealed a significant effect of subareas (Fig. 7G). Post-hoc test further showed that BN-HN in A1 L4 had the strongest significant GC links compared to A1 L2/3 and A2 L2/3, respectively. This result suggests that the proposed micronetwork is more well-represented in A1 L4 compared to the upper cortical layers.

Additionally, we explored the potential trend of decreased FC as the distances between BN and HN increases by performing Spearman correlation analysis. While A1 L4 and A2 L2/3 showed no significant trend as the distances changed, A1 L2/3 showed a small but significant trend of decrease of the mean values of top 5% GC links as the distance increased. Such result suggests that BNs in A1 L4 and A2 L2/3 does not preferably transfer information only to HNs in its neighborhood but also coordinate with HNs that are farther in the FOV.

Together, we showed that BNs can have significant but small directed excitatory influence onto HNs which is stronger in A1 L4 and weaker in L2/3 of A1 and A2. Specifically, the relatively stronger functional connectivity from BN to HN compared to randomized data could indicate that the linear inputs from BNs can serve as essential and meaningful building blocks for the nonlinear integration performed by HNs, ultimately shaping their sensitive responses to harmonic stacks. Such BN-HN connectivity is more well-represented in A1 L4.

### Parallel pathway contributions to response properties and nonlinear integration in ACtx neurons

ACtx receives parallel ascending inputs from the lemniscal and non-lemniscal pathways (Hackett et al., 2011; Saldeitis et al., 2014; Liu et al., 2019). These pathways shape the functional properties of ACtx neurons, with lemniscal input from the ventral medial geniculate body (MGBv) preferentially driving onset responses (Aitkin and Webster, 1972; Imig and Morel, 1983; Redies and Brandner, 1991; Hackett et al., 2011), while non-lemniscal input from the dorsal medial geniculate body (MGBd) is thought to preferentially driving offset responses (He, 2001; Liu et al., 2019). We investigated whether the temporal response properties of the three neuron types— harmonic-sensitive neurons (HNs), pure-tone neurons (PTNs), and broadly-tuned neurons (BNs)—aligned with lemniscal or non-lemniscal input patterns, focusing on their responses to sound onset and offset.

To characterize these temporal properties, we applied K-means clustering to the temporal response profiles of neurons, revealing distinct clusters dominated by onset or offset responses (Fig. 8A-B). We further quantified each neuron’s onset-offset bias using the offset bias index (OBI, OBI = (offset response – onset response) / (offset response + onset response)) (Liu et al., 2019) and compared OBIs across neuron populations (Fig. 8C-E). We observed differences in the OBI of sound-evoked responses of BNs from A1 L4 to A2 L2/3 and then to A1 L2/3 from onset- to offset-bias, with progressively greater onset bias for BNs in A2 L2/3 versus A1 L4. BNs in A1 L2/3 displayed significantly less onset bias than those in A1 L4 and A2 L2/3, with A2 L2/3 exhibiting an intermediate onset response (Fig. 8C). These findings suggest that BNs in A1 L4 may be more specialized in encoding the onset of sounds, while BNs in A1 L2/3 showed less preference for the onset of sounds. In contrast, OBIs for HNs were similar across areas (Fig. 8D), which indicates the similarity between harmonic-responsive neurons in A1 L4, A1 L2/3, and A2 L2/3 in detecting the initiation and termination of the harmonic stacks. The OBIs of PTNs in A2 L2/3 exhibited a significantly higher onset bias than A1 L4 (Fig. 8E), consistent with previous findings that A2 L2/3 neurons are more onset-biased in response to pure tones (Liu et al., 2019). We next analyzed OBIs of HNs to harmonic stacks of varied complexity (Fig. 8F) and found that OBIs of HNs were largely independent of both subareas and sound complexity. In summary, BNs show the most obvious subareal differences in onset-offset bias, PTNs in A1 L4 are significantly shifted to offset response compared to A2 L2/3, and HNs have homogeneous OBIs among the three subareas.

**Figure 8:**
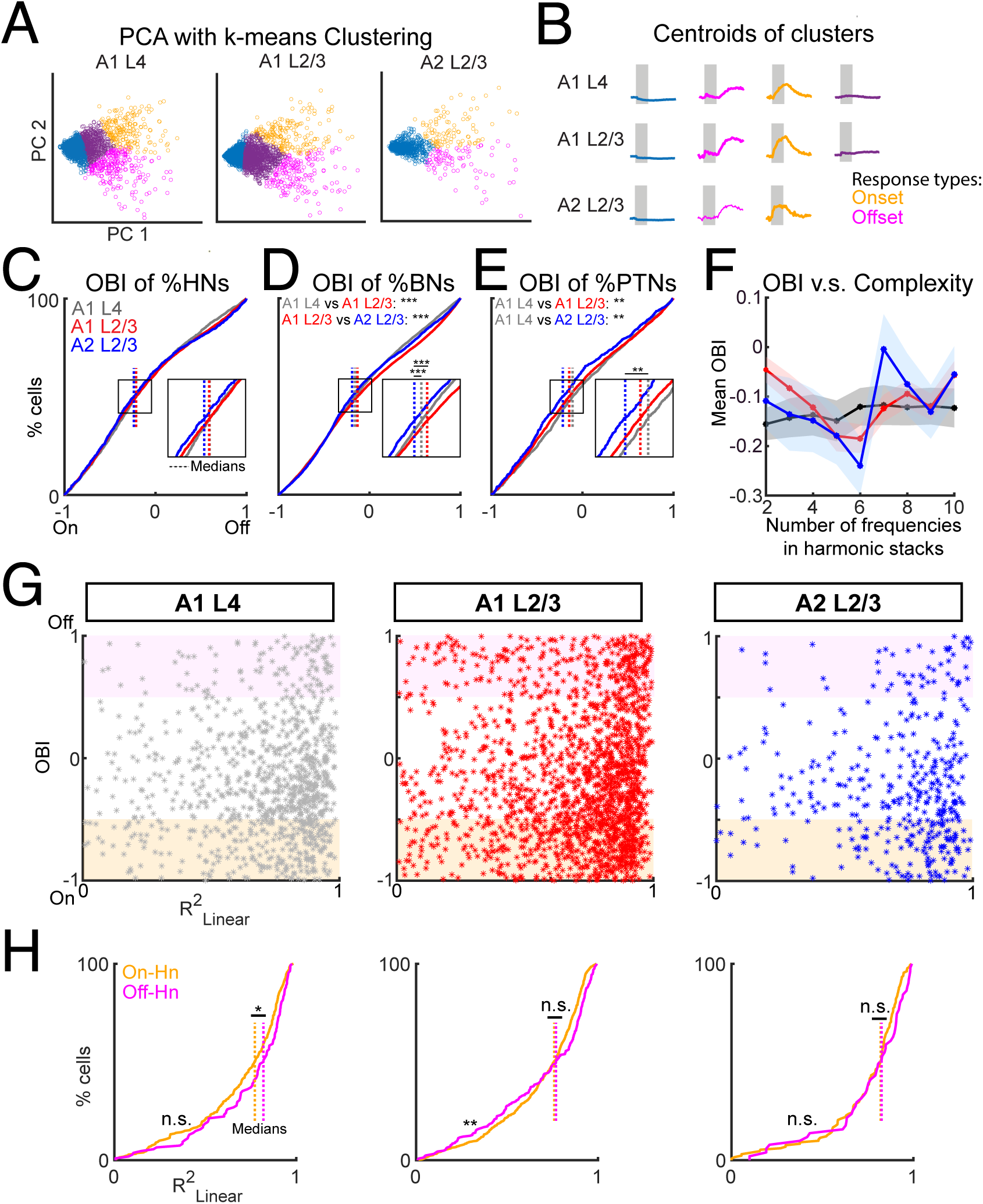
Neurons with offset-bias in A1 L4 has more linear integration. **A**: K-means clustering with varied numbers of clusters on the average sound-evoked response of HN to harmonic stacks. Each dot is the mean of 10 trials of sound-evoked response to one harmonic stack. Number of clusters is chosen using the elbow method for each subarea. In A1 L4, n = 2141 average traces from 1565 neurons. In A1 L2/3, n = 4811 traces from 3167 neurons. In A2 L2/3, n = 1002 average traces from 631 neurons. **B**: K-means clustering results that show distinct response types as offset response (magenta), onset response (orange). **C**: Cumulative probability of OBI of HNs. On the x-axis, -1 shows total onset biased, 1 shows total offset biased. Y-axis shows the cumulative probability of the OBIs. Dotted line shows median of OBIs in each subarea. Kruskal-Wallis test is performed due to the non-normal distribution of data. χ²(2) = 0.2177, p = 0.8969. Dunn’s post-hoc test with Bonferroni-corrected p-values on subareas: A1 L4 v.s. A1 L2/3, p = 1; A1 L2/3 v.s. A2 L2/3, p = 0.9539; A1 L4 v.s. A2 L2/3: p = 0.9755. Two-sample Kolmogorov-Smirnov test is performed on paired curves with post-hoc Bonferroni correction. Reported p-values are corrected. A1 L4 v.s. A1 L2/3, p = 0.2687; A1 L4 v.s. A2 L2/3, p = 0.2857; A1 L2/3 v.s. A2 L2/3, p = 1. **D**: Similar to C but for BNs in three subareas. Kruskal-Wallis test is performed due to the non-normal distribution of data. χ²(2) = 35.3030, p = 2.16E-8. Dunn’s post-hoc test with Bonferroni-corrected p-values on subareas: A1 L4 v.s. A1 L2/3, p = 1.45E-7; A1 L2/3 v.s. A2 L2/3, p = 9.75E-4; A1 L4 v.s. A2 L2/3: p = 0.9441. Cumulative distribution plot, two sample Kolmogorov-Smirnov test: A1 L4 v.s. A1 L2/3, p = 2.96E-17; A1 L4 v.s. A2 L2/3, p = 0.0734; A1 L2/3 v.s. A2 L2/3, p = 7.55E-5. **E**: Similar to C but for PTNs in three subareas. Kruskal-Wallis test is performed due to the non-normal distribution of data. χ²(2) = 10.8272, p = 0.0045. Dunn’s post-hoc test with Bonferroni-corrected p-values on subareas: A1 L4 v.s. A1 L2/3, p = 0.2101; A1 L2/3 v.s. A2 L2/3, p =0.0554; A1 L4 v.s. A2 L2/3: p = 0.0033. Cumulative distribution plot, two sample Kolmogorov-Smirnov test: A1 L4 v.s. A1 L2/3, p = 0.0093; A1 L4 v.s. A2 L2/3, p = 0.0752; A1 L2/3 v.s. A2 L2/3, p = 0.0034. **F**: OBI of HNs that respond to harmonic stacks with varied spectral complexity. Two-way ANOVA: Subareas: F(2,6144) = 0.73, p = 0.48; Harmonics: F(8,6144) = 1.96, p = 0.048; interaction: F(16, 6144) = 0.98, p = 0.48. Post-hoc t-test with corrected p-values show no significant among OBIs of HNs responding to different harmonic stacks. Solid line indicates the mean and the shaded area indicates the SEM. **G**: Scatter plots show linearity of response and OBI of every HN to the best harmonic sound in A1 L4, A1 L2/3 and A2 L2/3. The response linearity of HN with higher preference to onset-response (shaded by orange) or to higher offset-response (shaded by purple) are selected and plotted. **H**: Cumulative probability of R^2^ of linearity reconstruction on HNs response to harmonic sound by the response to pure-tone components in A1 L4, A1 L2/3 and A2 L2/3. Dotted line indicates the median of the distribution. Two-sample Kolmogorov-Smirnov test on the curves of on-HN and off-HN: A1 L4, p = 0.0667, A1 L2/3, p = 0.0091; A2 L2/3, p = 0.3650. Ranksum test on the medians of off-HN and on-HN: A1 L4, p = 0.0344; A1 L2/3, p = 0.2884; A2 L2/3, p = 0.3095

HNs exhibit nonlinear responses to simpler harmonic stacks, consistent with underlying mechanisms such as thresholding, where responses are only elicited when the summed input from multiple components exceeds a certain activation threshold, and rectification, where only positive or suprathreshold inputs drive significantly increased spiking activity, resulting in the increase of fluorescence change from the baseline in this study. Thus, to explore whether these nonlinear dynamics vary with response timing—particularly by extending response latency toward sound offset rather than onset—we analyzed the linearity of HNs with different OBIs. Specifically, we selected HNs with higher biases toward onset (shaded in beige) or offset (shaded in light purple) and examined their response linearity, respectively (Fig. 8G). Offset-biased HNs (off-HNs) in A1 L2/3 exhibit a more linear response profile compared to onset-biased HNs (on-HNs) (Fig. 8H), while HNs in A1 L4 showed a marginal trend toward this profile (p=0.0667), and HNs in A2 L2/3 showed no significant difference (p = 0.3650). These findings suggest that while HNs are similarly distributed across these cortical subregions, their response dynamics vary, with A1 HNs demonstrating a more linear onset response. This linearity may result from reduced inhibitory modulation and minimal positive feedback during spectral integration. Overall, this variation in response profiles across subregions highlights region-specific processing differences, potentially supporting distinct roles in auditory perception.

### Enhanced signal correlation and coordinated activity of harmonic neurons in A1 L4

Building upon our investigation of individual HN responses, we next aimed to characterize the collective activity of neurons sensitive to harmonic stacks but not to their pure tone components. While the proportion of HNs was similar across the three subareas (Fig. 4), prior widefield imaging studies revealed distinct activation patterns for tones and vocalizations in these regions (Calhoun et al., 2023). These differences may stem from varying reliability in how sound stimuli engage neuronal networks in each subarea. The sound-evoked response comprises a stimulus-driven component representing the overall response to the stimulus and a variable component that reflects the network’s activation pattern. The contributions of these components can be separated by calculating signal and noise correlations: signal correlations capture shared stimulus-driven responses, while noise correlations reflect functional connectivity between neurons (Averbeck et al., 2006; Averbeck and Lee, 2006; Cohen and Kohn, 2011; Winkowski and Kanold, 2013; Hazon et al., 2022). We calculated the signal correlation between HNs coactivated by each harmonic sound and compared across harmonic complexity and imaged subregions. Our results showed that increasing harmonic complexity did not significantly alter the signal correlation among HNs (Fig. 9A). Thus, more spectrally complex harmonic processing does not rely on recruiting additional neurons (Fig. 4D) or increasing synchronization of coactivated HNs as the harmonic becomes more spectrally broad (Fig. 9A). However, we observed that A1 L4 exhibited significantly higher signal correlations than A1 L2/3 and A2 L2/3, regardless of harmonic complexity (Fig. 9B). SCs were similar between A1 L2/3 and A2 L2/3. Thus, HNs in A1 L4 respond to harmonic stacks more synchronized than HNs in L2/3, potentially enabling A1 L4 to facilitate a more efficient, robust encoding of harmonic stacks through the heightened level of coordination. This high degree of coordination in A1 L4 may reflect a specialization for processing spectrally complex sounds, such as harmonic stacks, at an early stage of auditory processing.

**Figure 9:**
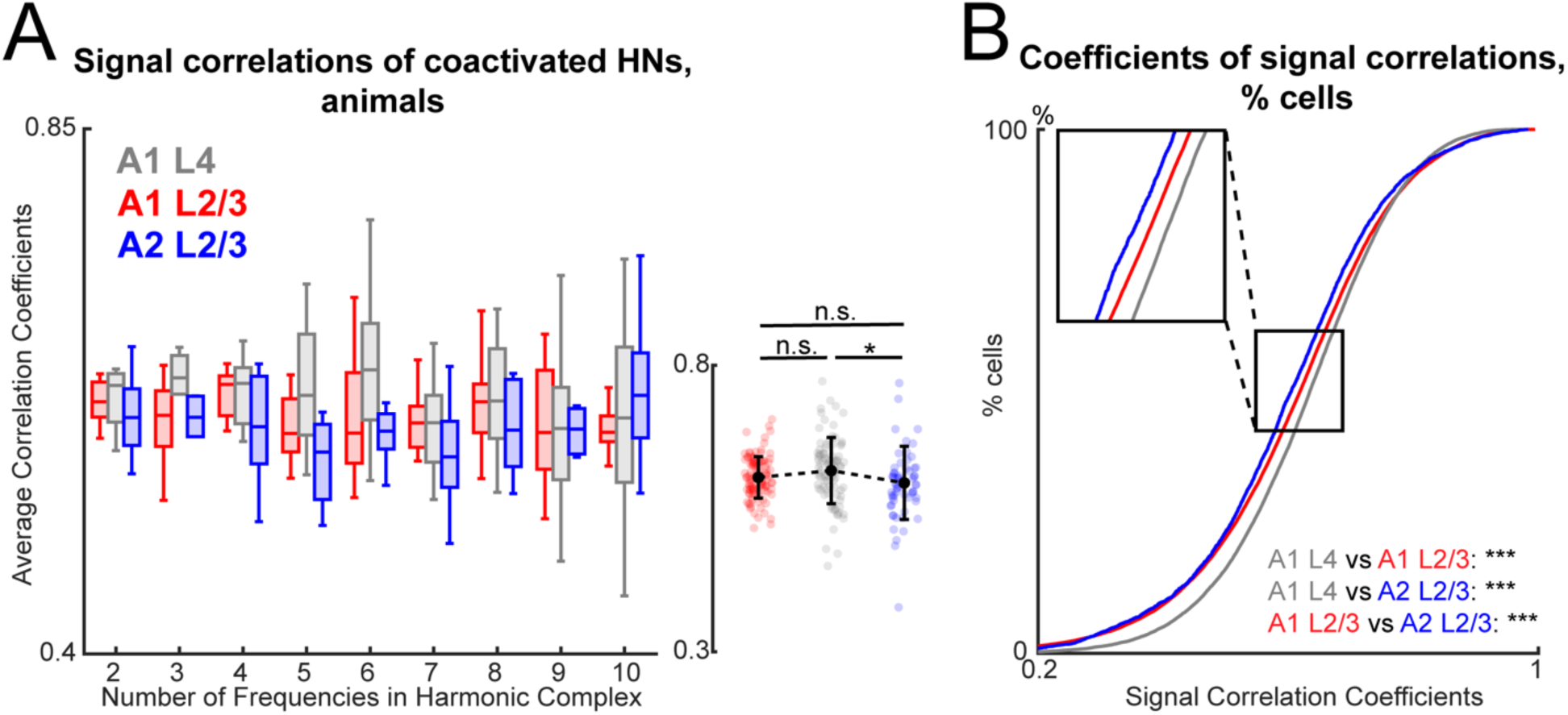
Hierarchical transformation of pair-wise signal correlation between coactivated HN pairs. **A**: Means of correlation coefficients (left) of pair-wise signal correlation between coactivated HNs responding to the activating harmonic stacks with varied numbers of components in A1 L4 (gray), A1 L2/3 (red) and A2 L2/3 (blue), as well as scatter plot (right) that shows the average signal correlation coefficients across different harmonic complexes. Each data point represents the mean coefficients of one subject. Two-way ANOVA on the left plot: Subareas, F(2,233) = 3.3990, p = 0.0351; harmonics, F(8, 233) = 1.5078, p = 0.1552; interactions, F(16, 233) = 0.8349, p = 0.6451. Post-hoc t-test with Tukey-Kramer correction on the scatter plots: A1 L4 v.s. A1 L2/3: p = 0.1552; A1 L4 v.s. A2 L2/3: p = 0.0280; A1 L2/3 v.s. A2 L2/3: p = 0.4656. **B**: Cumulative probability of pair-wise signal correlation coefficients in **(A)** between coactivated HNs in A1 L4 (gray), A1 L2/3 (red) and A2 L2/3 (blue). Two-sample Kolmogorov-Smirnov test on cumulative distribution curves with Bonferroni-corrected p-values: A1 L4 v.s. A1 L2/3: p = 1.1179E-53; A1 L4 v.s. A2 L2/3: p = 6.6488E-30; A1 L2/3 v.s. A2 L2/3: p = 5.0582E-6.

### Distinct functional connectivity and network sparsity of harmonic neurons in A1 L4

Noise correlations capture how fluctuations in neuronal activity, independent of stimulus-evoked response, are shared between pairs of neurons. High noise correlation indicates a high probability of shared connectivity between neurons, and thus noise correlation can serve as a proxy for measuring functional connectivity between neuron pairs. To further investigate network dynamics in harmonic processing, we examined the functional connectivity inferred by pairwise noise correlations between HNs and other neuron types, BNs, and PTNs.

To characterize stimulus-dependent network engagement, we compared noise correlations within HN pairs (HN-HN), between HNs and BNs (HN-BN), and between HNs and PTNs (HN-PTN) (Fig. 10). For HN-HN pairs, the ANOVA showed significant effects from both subareas and sound conditions on positive and negative noise correlations of HN-HN (Fig. 10B) despite that post-hoc test showed no significant differences among varying harmonic conditions on positive or negative correlations. We then examined whether there is a difference in the distribution of noise correlation coefficients between subareas and observed that HN-HN pairs in A1 L4 exhibited significantly lower positive and negative noise correlations compared to A1 L2/3 and A2 L2/3 (Fig. 10A), indicating that HNs in A1 L4 are less likely to form interconnected networks.

**Figure 10:**
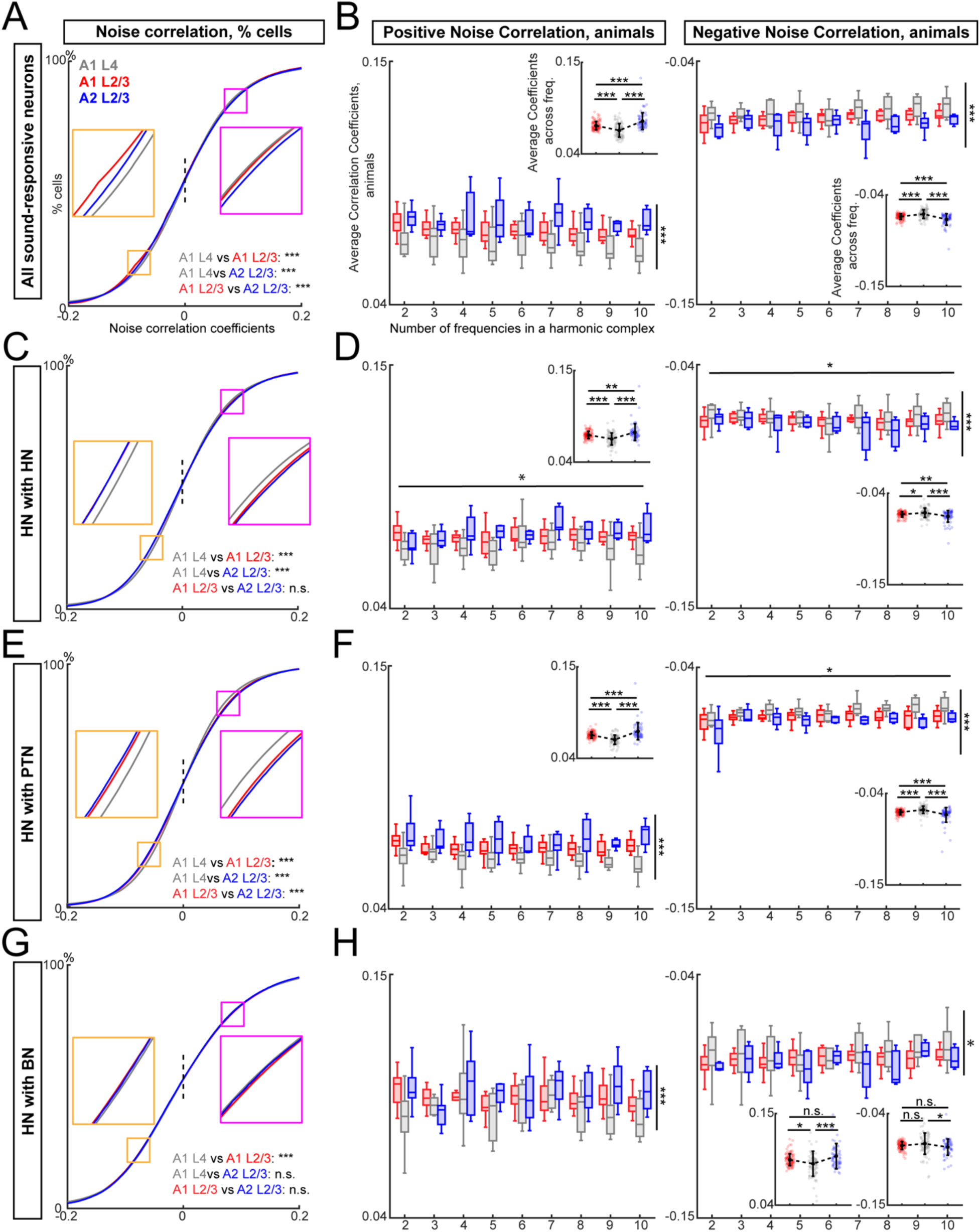
Noise correlation is the lowest in A1 L4, and similar in L2/3 of A1 and A2. **A:** Cumulative probability of the noise correlation coefficients of paired-neurons with significant facilitative response to any sound in three areas. Kolmogorov-Smirnov test for multiple groups with Bonferroni-corrected p-values: A1 L4 v.s. A1 L2/3: p = 4.52E-233; A1 L4 v.s. A2 L2/3: p = 2.93E-100; A1 L2/3 v.s. A2 L2/3: p = 1.78E-11. **B:** Positive (left) and negative (right) noise correlation coefficients between pairs of sound-responsive neurons in A1 L4 (gray), A1 L2/3 (red) or A2 L2/3 (blue) when different harmonic sound is present. Smaller box charts as insets shows the average noise correlation coefficients across sounds. Each data point is one animal in both plots. Positive correlation: Two-way ANOVA: Subareas, F (2,232) = 38.5750, p = 3.44E-15; harmonics, F (8, 232) = 0.6330, p = 0.7497; interactions, F (16, 232) = 0.5479, p = 0.9190. Post-hoc t-test with Tukey-Kramer-corrected p-values: A1 L4 v.s. A1 L2/3, p = 6.57E-7; A1 L4 v.s. A2 L2/3, p = 0; A1 L2/3 v.s. A2 L2/3, p = 3.58E-5. Negative correlation: Two-way ANOVA: Subareas, F (2,232) = 30.3274, p = 1.99E-12; harmonics, F (8, 232) = 1.1879, p = 0.3077; interactions, F (16, 232) = 0.8270, p = 0.6542. Post-hoc t-test with Tukey-Kramer-corrected p-values: A1 L4 v.s. A1 L2/3, p = 1.2E-4; A1 L4 v.s. A2 L2/3, p = 0; A1 L2/3 v.s. A2 L2/3, p = 3.54E-5. **C**: Cumulative probability of the noise correlation coefficients in three areas with presence of any harmonic sound. Kolmogorov-Smirnov test for multiple groups with Bonferroni-corrected p-values: A1 L4 v.s. A1 L2/3: p = 1.9493E-188; A1 L4 v.s. A2 L2/3: p = 7.3765E-48; A1 L2/3 v.s. A2 L2/3: p = 0.2990. **D**: Positive (left) and negative (right) noise correlation coefficients between pairs of HNs in A1 L4 (gray), A1 L2/3 (red) or A2 L2/3 (blue) when different harmonic sound is present. Smaller box charts as insets shows the average noise correlation coefficients across sounds. Each data point is one animal in both plots. Positive correlation: Two-way ANOVA: Subareas, F (2,234) = 24.8330, p = 1.6588E-10; harmonics, F (8, 234) = 2.0914, p = 0.0374; interactions, F (16, 234) = 1.2075, p = 0.2629. Post-hoc t-test with Tukey-Kramer-corrected p-values: A1 L4 v.s. A1 L2/3, p = 1.3102E-5; A1 L4 v.s. A2 L2/3, p = 0; A1 L2/3 v.s. A2 L2/3, p = 7.622E-3. Negative correlation: Two-way ANOVA: Subareas, F (2,234) = 13.0619, p = 4.1897E-6; harmonics, F (8, 234) = 2.1196, p = 0.0348; interactions, F (16, 234) = 1.0262, p = 0.4296. Post-hoc t-test with Tukey-Kramer-corrected p-values: A1 L4 v.s. A1 L2/3, p = 0.03349; A1 L4 v.s. A2 L2/3, p = 9.4061E-7; A1 L2/3 v.s. A2 L2/3, p = 6.533E-3. **E**: Similar to **(C)**, but between pairs of HN and PTN. Kolmogorov-Smirnov test for multiple groups with Tukey-Kramer-corrected p-values: A1 L4 v.s. A1 L2/3: p = 1.0340E-239; A1 L4 v.s. A2 L2/3: p = 1.1758E-77; A1 L2/3 v.s. A2 L2/3: p = 8.2345E-5. **F**: Similar to **(D)** but between pairs of HN and PTN. Positive correlation: Two-way ANOVA: Subareas, F (2,234) = 45.7036, p = 1.7549E-17; harmonics, F (8, 234) = 0.9572, p = 0.4703; interactions, F (16, 234) = 0.7702, p = 0.7185. Post-hoc t-test with Tukey-Kramer-corrected p-values: A1 L4 v.s. A1 L2/3, p = 0; A1 L4 v.s. A2 L2/3, p = 0; A1 L2/3 v.s. A2 L2/3, p = 3.0922E-Negative correlation: Two-way ANOVA: Subareas, F (2,234) = 23.9604, p = 3.4150E-10; harmonics, F (8, 234) = 2.2323, p = 0.0259; interactions, F (16, 234) = 0.9387, p = 0.5256. Post-hoc t-test with Tukey-Kramer-corrected p-values: A1 L4 v.s. A1 L2/3, p = 3.3942E-4; A1 L4 v.s. A2 L2/3, p = 0; A1 L2/3 v.s. A2 L2/3, p = 7.1524E-4. **G**: Similar to **(C)**, but between pairs of HN and BN. Kolmogorov-Smirnov test for multiple groups with Tukey-Kramer-corrected p-values: A1 L4 v.s. A1 L2/3: p = 6.6985E-8; A1 L4 v.s. A2 L2/3: p = 0.0818; A1 L2/3 v.s. A2 L2/3: p = 0.5404. **H**: Similar to **(D),** but between pairs of HN and BN. Positive correlation: Two-way ANOVA: Subareas, F (2,229) = 9.8325, p = 8.0033E-5; harmonics, F (8, 229) = 0.5160, p = 0.8439; interactions, F (16, 229) = 0.9143, p = 0.5535. Post-hoc t-test with Tukey-Kramer-corrected p-values: A1 L4 v.s. A1 L2/3, p = 0.0230; A1 L4 v.s. A2 L2/3, p = 3.2149E-5; A1 L2/3 v.s. A2 L2/3, p = 0.06987. Negative correlation: Two-way ANOVA: Subareas, F (2, 229) = 3.6435, p = 0.0277; harmonics, F (8, 229) = 0.6496, p = 0.7354; interactions, F (16, 229) = 0.4749, p = 0.9573. Post-hoc t-test with Tukey-Kramer-corrected p-values: A1 L4 v.s. A1 L2/3, p = 0.2106; A1 L4 v.s. A2 L2/3, p = 0.0211; A1 L2/3 v.s. A2 L2/3, p = 0.4225. Whiskers in larger box charts represent SEM. Whiskers in smaller box charts within the larger charts represent standard deviation.

Similarly, we examined HN-PTN and HN-BN noise correlations. For HN-PTN pairs, the ANOVA revealed significant effects of subareas and sound condition on negative noise correlations but not on positive noise correlation (Fig. 10D). Post hoc analysis showed HN-PTN noise correlations were lowest in A1 L4 compared to A1 L2/3, and A2 L2/3, which had the highest noise correlation among the three areas (Fig. 10C). For HN-BN pairs, the harmonic sound condition had no significant effect, but subarea remained a key factor, with A1 L4 displaying significantly lower noise correlations compared to A1 L2/3 (Fig. 10F). No significant differences were found between A1 L4 and A2 L2/3, or between A1 L2/3 and A2 L2/3 (Fig. 10E).

Together, our analysis of noise correlations among neuron types across subareas suggests that HNs in A1 L4 are more likely to receive distinct synaptic inputs, forming sparser and more selective networks than HNs in L2/3, which exhibit stronger noise correlations. This network sparsity in A1 L4 remained consistent despite increasing spectral complexity of harmonic stacks, supporting the idea that the ACtx encodes sounds sparsely and selectively adapts to varying spectral complexity.

## Discussion

We investigated the population coding of spectrally simple and complex harmonic stacks in different auditory cortical subfields. We find harmonic-sensitive neurons (HNs), that respond to harmonic stacks but not to single frequency. We find that multiple subareas, including A1 L4, A1 L2/3 and A2 L2/3, in the ACtx contain HNs, and that the fraction of HNs is similar across A1 and A2. Thus, harmonic sensitivity is already present in A1 L4 and is not a unique feature of A1 L2/3 and A2 L2/3. HNs show sensitivity to particular stacks of harmonic frequencies and are characterized by nonlinear integration of the component frequencies.

Simple sounds, such as pure tones or harmonic stacks of few frequencies, serve as the fundamental building blocks of more intricate stimuli, such as harmonic stacks of more than five frequencies found in speech. Although previous studies have examined the neural representation of specific sound features in the ACtx of humans, rats, and mice (Lewis et al., 2009; Okada et al., 2010; Carruthers et al., 2015; de Heer et al., 2017; O’Sullivan et al., 2019; Staib and Fruhholz, 2023), the question remains of if the spectral integration of harmonic stacks with simpler to more complex structures varies across ACtx layers and subareas. Our results show that A1 L4, A1 L2/3, and A2 L2/3 contain neurons that are sensitive to harmonic stacks with no significant response to any pure tone, and that proportions of HNs are largely similar across subareas while their functional connectivity differs. Thus, HNs seem to be independently assembled in multiple ACtx areas but form distinct networks.

Intrinsic imaging has suggested that A2 L2/3 preferentially activated by harmonic (Kline et al., 2021). In contrast, we found that the proportion of neurons activated by harmonic stacks was similar across A1 L4, A1 L2/3, and A2 L2/3, regardless of number of harmonic frequencies in the stack or the sex of the mice. The differences between our study and the prior study (Kline et al., 2021) likely lie in the imaging specificity, duration of sounds, as well as mouse lines. Instead of intrinsic imaging with low spatial resolution, we used in vivo two-photon imaging and electrophysiologywith single cell resolution. Instead of short duration sounds (100–300 ms), we utilized sound with longer duration (1000 ms). Certain proportions of neurons may be sensitive to the duration of the sound (Theunissen et al., 2000; Buonomano and Maass, 2009), or require longer stimuli to trigger the significant changes in calcium traces. The prior study utilized C57Bl/6 mice and there are also sex-dependent age-related changes in hearing (Shilling-Scrivo et al., 2021, 2022), and thus the differences could also be due to C57Bl/6 mice having early-onset high-frequency hearing loss (Ison et al., 2007; Jendrichovsky et al., 2024). In contrast, we use mice that retain good high-frequency hearing across age. Moreover, given the behavioral importance of natural stimuli containing harmonic stacks, the sensory experience of animals could shape the responses in ACtx, thus differences in the rearing environment (Chang and Merzenich, 2003; Sanes and Bao, 2009; Homma et al., 2020; Chang and Kanold, 2021) could underlie the observed differences.

Among HNs in three imaged auditory subfields, we showed that spectral shift of frequency disrupted their response to the non-shifted harmonic two-tone stacks. In this study, the spectral shift was applied only in the downward direction, disrupting harmonicity and narrowing spectral bandwidth. As a result, the diminished responses could reflect sensitivity to either the altered frequency relationship or changes in bandwidth. Nonetheless, the consistent reduction in activity across regions suggests that these neurons respond preferentially to specific frequency combinations with harmonic structure. Future studies incorporating upward or bidirectional shifts could be utilized to distinguish the relative contributions of harmonicity and spectral bandwidth.

The harmonic-sensitive response of HNs can be explained by their nonlinear integration of component responses. HN responses become more linear with increased number of harmonic frequencies, suggesting that broader frequency integration is associated with increased linearity. Consistent with studies reporting supra-linear and sublinear integration in A1 L2/3 and A2 L2/3 for harmonic representation (Kline et al., 2023), our results provided insights of how the linearity of spectral integration can change depending on the spectral contents. We observed that neurons’ responses to harmonic stacks with fewer frequencies were highly nonlinear in A1 L4, A1 L2/3 and A2 L2/3 (Fig. 6). Such nonlinear integration emerges earlier than L2/3 and might also be observable in auditory thalamus. This nonlinearity in spectral integration diminished as the number of harmonic frequencies increased in a stack in all imaged subfields. This is potentially due to reduced thresholding in the local network as the number of harmonic frequencies increases. It is also possible that the representation of simpler and more complex harmonic stacks are through the two different pathways, linear and nonlinear pathways, emerging from the cochlear nucleus (Yu and Young, 2000). These findings underscore the dynamic nature of spectral integration in the ACtx, adapting to the varied complexity of auditory stimuli.

As we characterized the nonlinear spectral integration of frequencies by HNs, we proposed a micronetwork model, delineating potential structural and functional connectivity that might underly the harmonic-sensitive response tuning of HNs. (Fig. 7). Notably, broadly-tuned neurons (BNs) appear critical for establishing harmonic-sensitive tuning: they display lower sensitivity for particular harmonic stacks, and their pure-tone response reconstructs harmonic responses with higher linearity. This suggest that BNs may generalize across frequencies and facilitate the harmonic sensitivity in HNs. Granger causality (GC) analysis revealed a small but significantly stronger directed influence from BNs to HNs in A1 L4 compared to L2/3 of A1 and A2. This result suggests that the underlying network may not be confined to A1 L4, but rather originate earlier in the auditory pathway which is known to shape and refine representations of complex acoustics (Patterson et al., 2002; Nelken, 2008; Bartlett, 2013). Moreover, signal correlations among A1 L2/3 neurons are patchy and stimulus-dependent, suggesting upstream convergence and nonlinear integration (Jendrichovsky et al., 2025). Given the spatial encoding of single frequencies in mammalian cochlea (Dallos, 1996) and the sparse representation of vocalization features observed in cortex across multiple species (Hromadka et al., 2008; Bandyopadhyay et al., 2010; Bowen et al., 2020; Montes-Lourido et al., 2021), we speculate that the BN-> HN micronetwork spans multiple brain regions and enables the special tuning profiles of HNs in the auditory cortex.

To explore the sources of nonlinearity, we examined whether nonlinear neurons might show preference to the sound onset or offset, which are essential for auditory scene analysis (Bregman, 1994). Our study showed that BNs and PTNs are more onset-biased in A1 L4 compared to A2 L2/3. Specifically, BNs in A2 L2/3 also show significantly more onset bias compared to those in A1 L2/3. Which is consistent with our previous finding that A1 neurons were more off-set biased to pure tones compared to those in A2 (Liu et al., 2019), without evaluating the responses to harmonic stacks; thus, neurons in the previous study likely were a combination of PTNs and BNs in this study. As A1 L2/3 receives inputs from both A1 L4 and A2 L2/3, it may be an integrative hub for processing diverse spectral information in sustained auditory stimuli. The observed hierarchy across layers and regions likely reflects specialized roles in temporal processing. Notably, in A1 L2/3, onset-biased HNs tended to be more linear, while offset-biased HNs were more nonlinear. These results suggest that the onset pathway may encode spectrally complex sounds more linearly, with increased nonlinearity in the non-lemniscal offset pathway.

Coactivated HNs in A1 L4 exhibit the highest signal correlations compared to those in A1 L2/3 and A2 L2/3. This result contradicts to the hypothesized hierarchy of harmonic processing from A1 L4 to A1 L2/3 and finally to A2 L2/3. An alternative explanation for this non-hierarchical transformation is that neurons in L2/3 may engage in a more distributed representation of harmonic features, resulting in a lower synchronized sound-evoked response compared to A1 L4. This could indicate that as sound information progresses from L4 to L2/3, it undergoes further abstraction, leading to reduced coordination among L2/3 neurons. This distinction suggests that A1 L4 may play a foundational role in the initial encoding of harmonic structure, while L2/3 may contribute to higher-order processing of sound features. The neuronal network involving all sound-evoked neurons in A1 L4 was the sparsest, showing the lowest noise correlation. This is consistent with our previous study which found pure-tone responsive neurons in A1 L4 showed higher signal correlations and lower noise correlations than those in A1 L2/3 (Winkowski and Kanold, 2013). Here A1 L4 HNs also showed low noise correlations with pure-tone and broadly-tuned neurons compared to L2/3 suggesting that A1 L4 neuron populations show less of the shared synaptic inputs regardless of their sound-response profiles.

Our results show that harmonically sensitive neurons are ubiquitous in A1 and A2, indicating the importance of harmonic integration for auditory processing. Thus, already, A1 L4 functions as a center for integrating spectral information for complex sounds, supporting robust encoding through coordinated but sparsely connected networks.

## Acknowledgements & Contributions

YC and POK designed research. YC performed imaging experiments and analysis. YG performed imaging analysis. CC and YC performed electrophysiological experiments and analysis. YC and POK drafted paper. All authors edited the manuscript. Supported by NIH RO1DC017785 and NIH RO1DC009607 (POK).

